# Gentle label-free nonlinear optical imaging relaxes linear-absorption-mediated triplet

**DOI:** 10.1101/2023.10.09.561579

**Authors:** Geng Wang, Lianhuang Li, Janet E. Sorrells, Jianxin Chen, Haohua Tu

**Affiliations:** Department of Electrical and Computer Engineering, University of Illinois at Urbana-Champaign, Urbana, IL, 61801, USA; Beckman Institute for Advanced Science and Technology, University of Illinois at Urbana-Champaign, Urbana, IL, 61801, USA; Key Laboratory of OptoElectronic Science and Technology for Medicine of Ministry of Education, Fujian Provincial Key Laboratory of Photonics Technology, Fujian Normal University, Fuzhou, 350007, China

**Author notes:** These authors contributed equally to this work (Geng Wang, Lianhuang Li).

## Abstract

Sample health is critical for live-cell fluorescence microscopy and has promoted light-sheet microscopy that restricts its ultraviolet-visible excitation to one plane inside a three-dimensional sample. It is thus intriguing that laser-scanning nonlinear optical microscopy, which similarly restricts its near-infrared excitation, has not broadly enabled gentle label-free molecular imaging. We hypothesize that intense near-infrared excitation induces phototoxicity via linear absorption of intrinsic biomolecules with subsequent triplet buildup, rather than the commonly assumed mechanism of nonlinear absorption. Using a reproducible phototoxicity assay based on the time-lapse elevation of auto-fluorescence (hyper-fluorescence) from a homogeneous tissue model (chicken breast), we provide strong evidence supporting this hypothesis. Our study justifies a simple imaging technique, e.g., rapidly scanned sub-80-fs excitation with full triplet-relaxation, to mitigate this ubiquitous linear-absorption-mediated phototoxicity independent of sample types. The corresponding label-free imaging can track freely moving *C. elegans* in real-time at an irradiance up to one-half of water optical breakdown.

## Introduction

Due to plausible artifacts from light itself^**1**^, live-cell fluorescence imaging has increasingly emphasized sample health over metrics such as signal-to-noise ratio (SNR) and spatiotemporal resolution^**2**^. The classic mechanism of phototoxicity (Fig. 1a) attributes the toxicity to the excited singlet and triplet states of extrinsic (labeling) UV-visible-absorbing fluorophores that induce reactive oxygen species (ROS)^**2**,**3**^. Thus, wide-field planar excitation that restricts the excitation to a focal plane (e.g. light-sheet microscopy) has gained popularity to avoid out-of-focus ROS production in standard wide-field or laser-scanning fluorescence microscopy^**3**,**4**^. Another technique is to excite the labeling fluorophores at the long-wavelength end of UV-visible excitation (300-650 nm), which mitigates the phototoxicity from intrinsic ROS-generating photosensitizers^**5**^ (Fig. 1a) but limits the choices to fluorescently label the sample. By avoiding this critical limitation from popular continuous-wavelength excitation, near-infrared (NIR) extension (700-1300 nm) with laser-scanning ultrashort (<10 ps) pulses has recovered the planar excitation and efficiently excited the extrinsic fluorophores via nonlinear absorption^**6**^ (Fig. 1b).

**Fig. 1.**
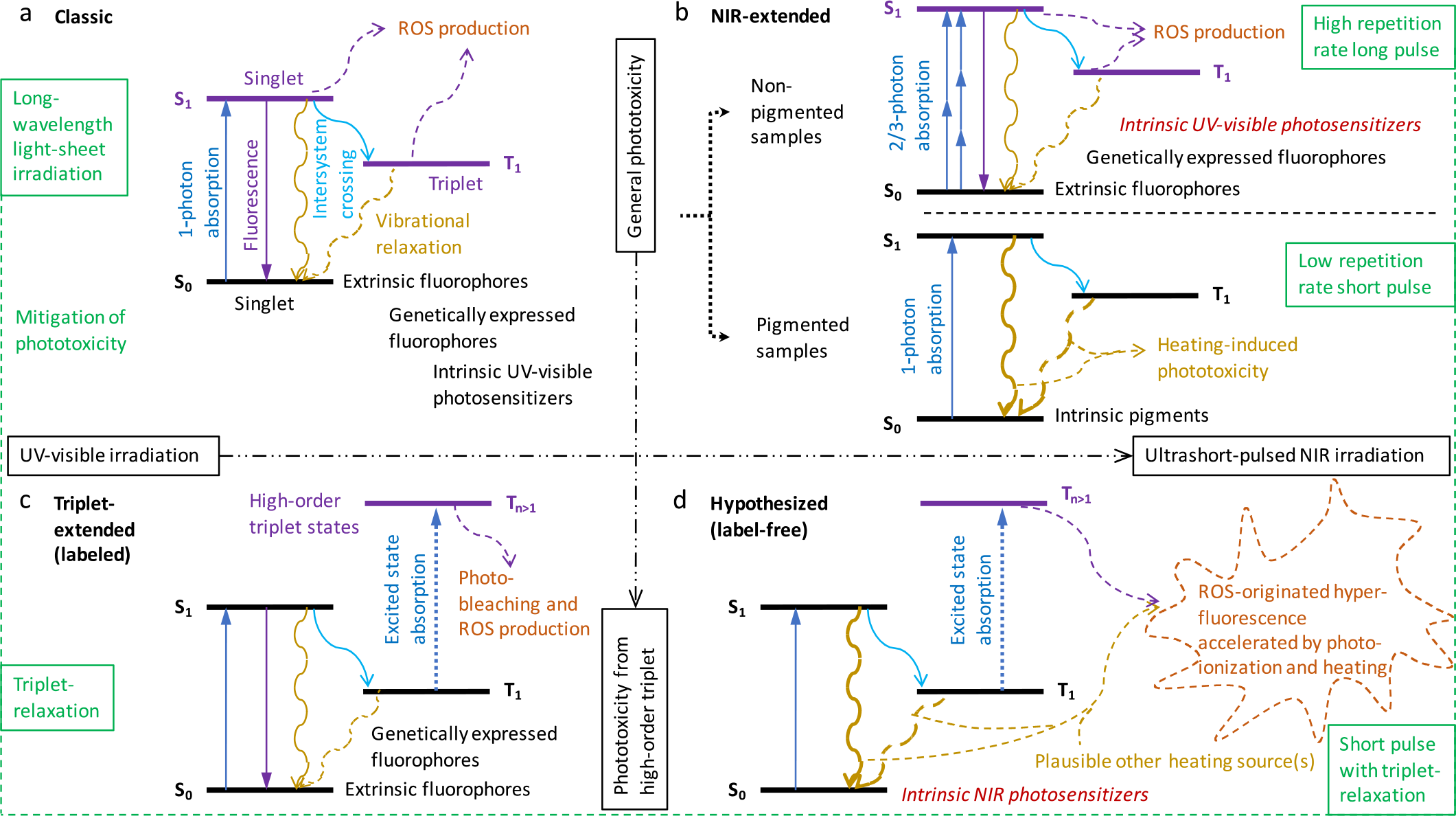
Mechanisms of phototoxicity based on Jablonski diagram. **a** Classic mechanism with UV-visible excitation. **b** NIR-extended mechanism with scanning ultrashort-pulsed NIR excitation at the focus of a microscope objective. **c** Triplet-extended mechanism with UV-visible excitation of labeling fluorophores. **d** Hypothesized mechanism at phototoxicty threshold with the same NIR excitation but without the labeling. One-photon absorption of certain ubiquitous chromophore followed by efficient inter-crossing to a first-order triplet state results in high-order triplet-induced heating-accelerated white hyper-fluorescence (WHF).

However, raster scanned ultrashort-pulsed NIR excitation at the focus of a high numerical aperture (NA) microscope objective has either facilitated linear-absorption-mediated heating toxicity in pigmented samples, which should be mitigated by low repetition rate short pulse^**7**^, or commonly assumed nonlinear-absorption-mediated phototoxicity in non-pigmented samples^**8**,**9**^, which should be mitigated by high repetition rate long pulse^**10**,**11**^. Because the two are differentiated rather arbitrarily based on often unavailable NIR absorption property of the sample^**12**^, no universal technique exists for both cases (Fig. 1b). Also, the related NIR-extended mechanism favors a light-dose (fluence) threshold for phototoxicity^**9**,**13**^ over an irradiance threshold (typically proportional to average power or pulse energy), which is inconsistent with empirical experiences^**8**,**14**^ (Supplementary Table 1). Moreover, this mechanism offers no satisfactory explanation on why the phototoxicity decreases with increased speed of fast-axis scanning, even though other parameters are comparable (Supplementary Table 2).

We aim to overcome these deficiencies in knowledge based on another extension of the classic mechanism, i.e., triplet-extended mechanism of photobleaching-related phototoxicity^**2**,**15**^ from the labeling agents of extrinsic fluorophores and genetically expressed fluorophores^**16**^ (Fig. 1c). This mechanism offers a satisfactory explanation (i.e., triplet-relaxation) on why a fast-scanning speed effectively mitigates the phototoxicity in both confocal fluorescence microscopy^**17**^ and stimulated emission depletion microscopy^**18**,**19**^. Specifically, high-order triplet states produced by excited state absorption^**16**^, rather than first-order triplet/singlet states and high-order singlet states, dominate the observed UV-visible phototoxicity from bioassays^**17**,**19**^ (Fig. 1c). Considering the similar dependence of the NIR phototoxicity on the scanning speed (Supplementary Table 2), we hypothesize that the corresponding NIR extension would follow a similar mechanism (Fig. 1d) in label-free (clinically permissible) nonlinear optical imaging (Supplementary Table 3). The focus on the label-free scenario in this study will ultimately guide more complicated scenarios with diverse labeling fluorophores.

Our hypothesized mechanism asserts that linear absorption of intrinsic NIR photosensitizers mediates the NIR phototoxicity in unlabeled samples (Fig. 1d), rather than the commonly assumed nonlinear absorption of intrinsic UV-visible photosensitizers^**8**^ (Fig. 1b) such as NAD(P)H, flavins, and porphyrins^**14**^. This is plausible because in a broad (non-imaging) context, the existence of intrinsic NIR photosensitizers has been demonstrated in *E. coli* inside an optical trap^**20**^ and cultured cells under phototherapy^**21**^ capable of ROS production^**22**^. Another unusual feature is the enhancement of high-order triplet-mediated ROS production by linear absorption-induced heating (Fig. 1d) like that in pigmented samples such as skin and retina (Fig. 1b). This is inspired by the heating-accelerated phototoxicity in excessive photobiomodulation^**23**^, and as shown below, can reconcile various contradictory views on the NIR phototoxicity (Supplementary Table 1). Surprisingly, all evidence that favor the nonlinear-absorption-mediated phototoxcity in unlabeled samples (Fig. 1b, Supplementary Table 4) can be alternatively interpreted by the hypothesized mechanism to reconcile with those that favor the linear-absorption-mediated phototoxicity in non-pigmented samples (Supplementary Tables 5,6).

## Results

### Chicken breast model quantifies homogeneous hyper-fluorescence

Our study is motivated by the intrinsic indicator of phototoxicity specific to nonlinear optical microscopy, i.e., elevated auto-fluorescence during time-lapse imaging of diverse (live) cell/tissue specimens^**24**^. This effect has been observed by different groups with widely varied excitation-scanning parameters and samples, resulting in different terminologies such as “white flashes”, “flickering/broadband luminescence”, “fluorescent scar/lesion”, “photo-modulation”, “photo-enhancement”, and “hyper-fluorescence” (Supplementary Table 5). Despite the established functional link to impaired cell cloning^**8**^ and ROS/apoptosis^**25**^, these different terminologies may have hindered a general understanding of the same underlying phenomenon not observable by linear optical microscopy. We adopt “white hyper-fluorescence” (WHF) throughout this paper to emphasize its broadband emission and inline (built-in) indication of phototoxicity^**26**^ in unlabeled biological samples^**24**^.

We employed simultaneous label-free autofluorescence-multiharmonic (SLAM) microscopy in time-lapse imaging of live cells (Fig. 2a) and *ex vivo* mouse kidney tissue (Fig. 2b), with four simultaneously acquired molecular contrasts of two-/three-photon auto-fluorescence (2PAF/3PAF) and second-/third-harmonic generation (SHG/THG)^**27**^. We built a portable SLAM microscope (pSLAM) with more flexible excitation-scanning parameters^**28**^ and an extended version of SLAM microscope (eSLAM) with a faster imaging speed^**29**^. The point-spreading heterogeneous WHF detected by eSLAM across 2PAF-3PAF detection spectrum of 420-640 nm (Fig. 2a,b, arrowheads; Extended Data Table 1; Supplementary Videos 1,2) was not amenable for quantification. In contrast, under an illumination of pSLAM (Extended Data Table 2, spatiotemporal bin-10) except for a higher average power, we observed the emergence of homogeneous WHF in chicken breast tissue across a large area of field-of-view (FOV) (Fig. 2c, broken cycle; Supplementary Video 3) before the occurrence of similar heterogeneous WHF (Fig. 2c, arrowheads) and subsequent bubble formation (Fig. 2c, stars). Lowering the power avoided the heterogeneous WHF in time-lapse imaging, during which the homogeneous WHF underwent linear growth initially (Fig. 2d, arrowed lines) rather than a decrease via photo-bleaching, until a power threshold at WHF onset was reached (14 mW in Extended Data Fig. 1a). The homogeneous WHF attains a spatial distribution that approximates the illumination field measured in a fluorophore solution^**29**^, indicating the field strength-dependent phototoxicity. The increased THG signal most likely arises from the blue edge of WHF, while the corresponding increase in SHG signal may be canceled by a decrease due to thermal denaturing^**30**^ (Fig. 2c,d, bottom panels).

**Fig. 2.**
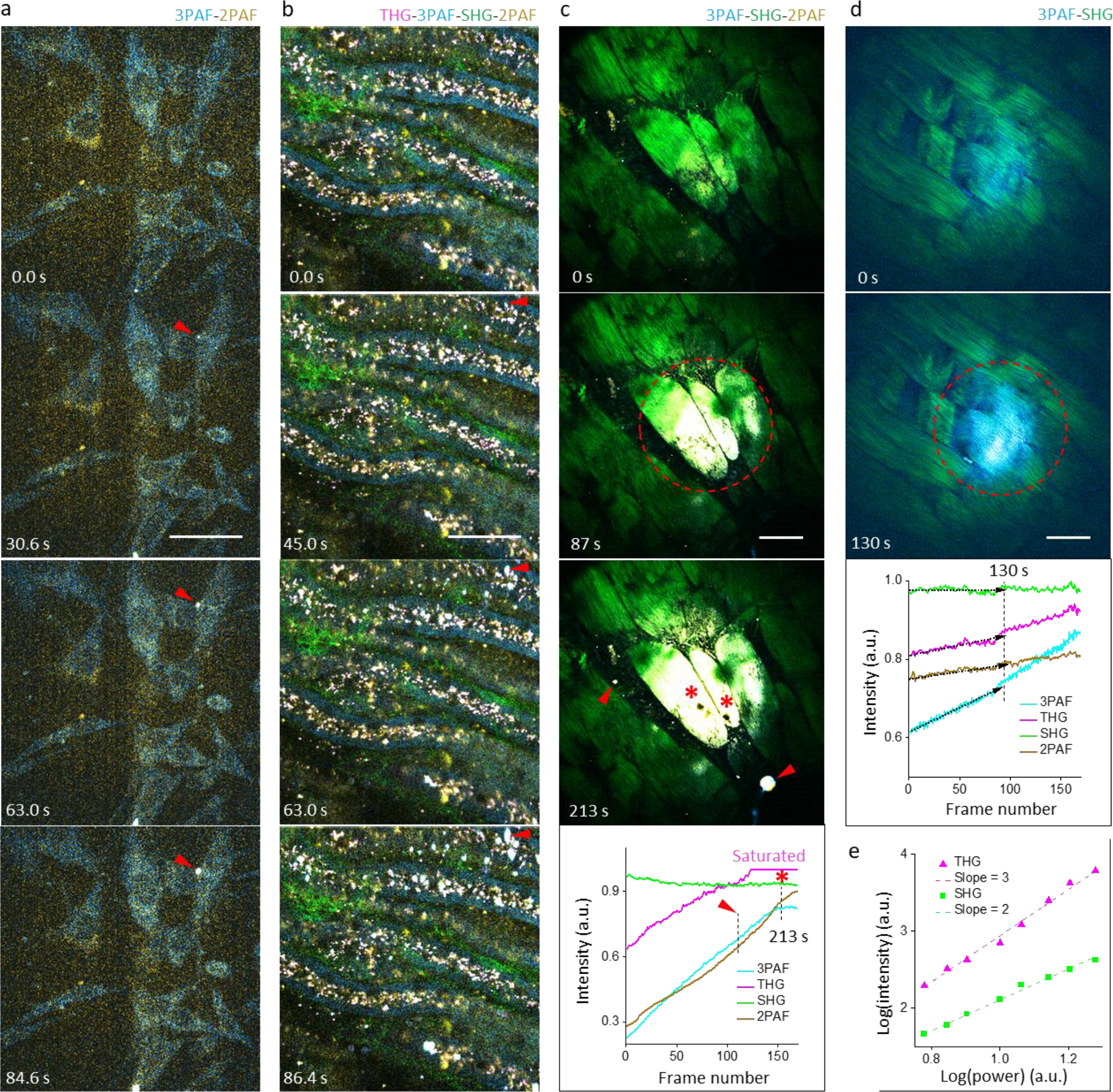
Phototoxicity observed during time-lapse SLAM-based imaging. **a** Cultured hamster kidney cells imaged by eSLAM at a frame rate of 0.56 Hz, showing heterogeneous “white” heterogeneous WHF due to simultaneous increase of 3PAF/cyan and 2PAF/yellow signals (arrowheads). **b** *ex vivo* mouse kidney tissue imaged by eSLAM at a frame rate of 0.56 Hz, showing similar WHF (arrowheads). **c** chicken breast imaged by pSLAM (25 mW, spatiotemporal bin-10) at a frame rate of 0.7 Hz, showing emergence of homogeneous WHF (broken cycle) followed by the heterogeneous WHF (arrowheads) and subsequent bubble formation (stars); bottom panel shows integrated signals over one frame versus frame number during time-lapse imaging. **d** chicken breast imaged by pSLAM at a lower power (15 mW, spatiotemporal bin-10), showing only the homogeneous WHF (broken cycle); bottom panel shows the integrated signals with initial linear growth (arrowed lines). **e** integrated THG/SHG signals versus power (baseline illumination) in different FOVs that follow power-3/power-2 law according to the photon order of nonlinear optical processes. Scale bar: 50 μm.

These results are in sharp contrast to the reported heterogeneous WHF where no linear growth rate and correlation to the illumination field has been established (Fig. 2a,b; Supplementary Table 5). In contrast to the heterogeneous WHF and bubble formation, the homogeneous WHF is more suitable for quantification due to the uniform morphology, independence on dosage, and linear growth at an early stage of phototoxicity. Under another illumination (Extended Data Table 2, baseline) except for a variable power, different FOVs in one sample of chicken breast at one controlled imaging depth (15±5 μm fixed) largely follows the power laws of nonlinear signals, with a small error of <20% (Fig. 2e). We therefore selected chicken breast as a readily available model to reproducibly quantify the homogeneous WHF under different illuminations.

### Linear rather than nonlinear absorption induces hyper-fluorescence

To evaluate the effect of pulse width^**8,26**^ on 2PAF/3PAF linear growth rates (arbitrary unit per pulse), we compared two illuminations 10% above the corresponding WHF power thresholds (Extended Data Table 2, baseline vs. chirped), which produced consistent data across different testing FOVs (Fig. 3a, top; Extended Data Fig. 2). By taking account of the dual role of excitation pulses to induce and detect WHF, the observed growth rates associated with the chirped illumination (Extended Data Fig. 3, left) can be predicted from those associated with the baseline illumination according to an assumed phototoxicity photon order of either 1 or 2. Because the former is in quantitative agreement with experimental data (Fig. 3a, bottom), the observed WHF is induced by linear absorption (Fig. 1d).

**Fig. 3.**
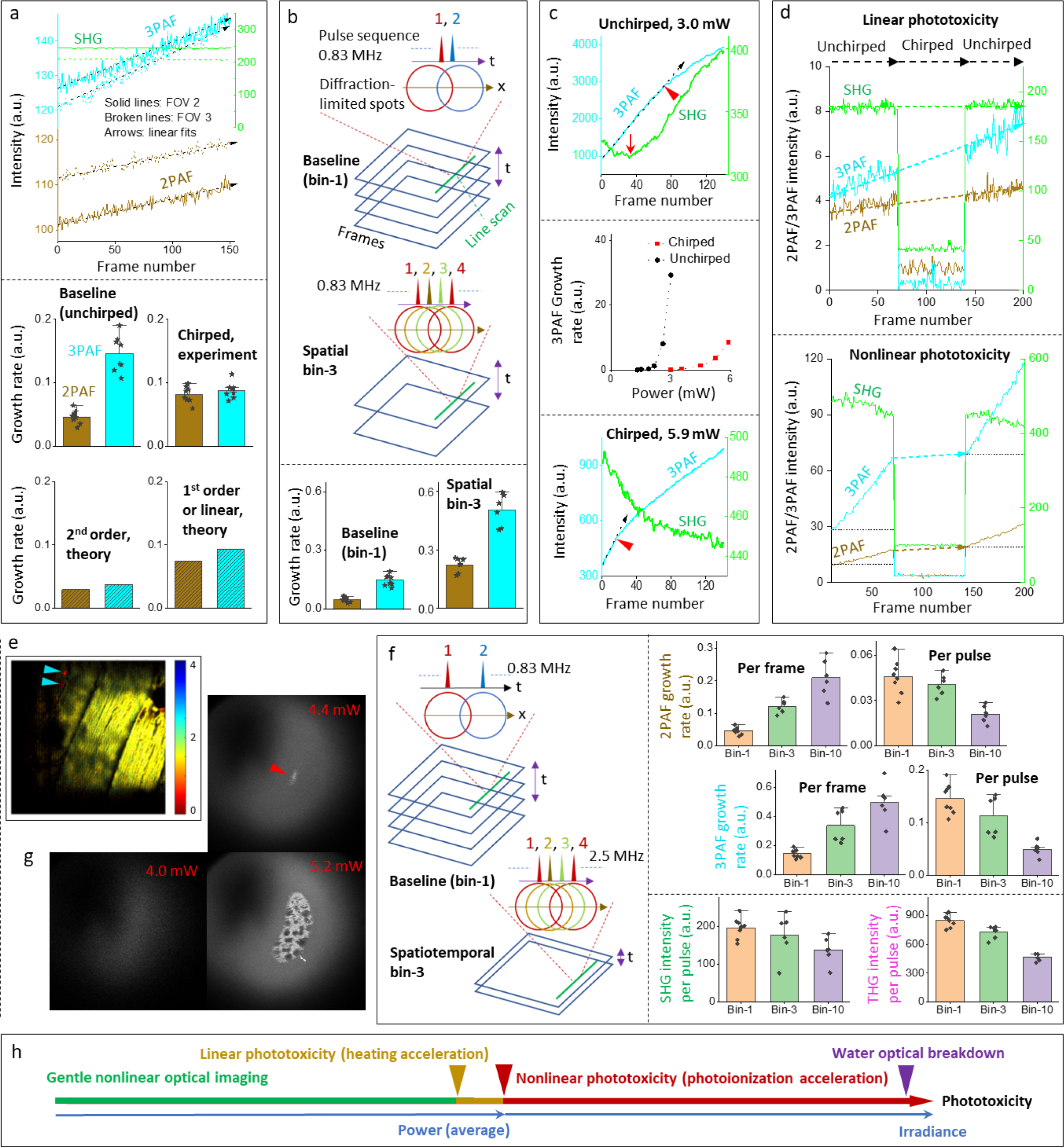
Characteristic features of homogeneous WHF-revealed phototoxicity. **a** (Top) Reproducible 2PAF and 3PAF growth rates under the baseline illumination of time-lapse pSLAM imaging in two different FOVs of chicken breast, despite their difference in absolute intensity integrated over one frame; (Bottom) observed 2PAF and 3PAF growth rates under pSLAM baseline/unchirped illumination (upper left) and chirped illumination (upper right), along with calculated rates of the latter according to 2^nd^ order phototoxicity (lower left) and 1^st^ order or linear phototoxicity (lower right). **b** (Top) Illustration of spatial bin-3 illumination in comparison to baseline/bin-1 illumination; (Bottom) comparison of 2PAF and 3PAF growth rates according to bin-1 (left) and bin-3 pSLAM illumination (right). **c** (Top) High WHF-revealed phototoxicity but late bubble formation (arrowhead) at a high pSLAM irradiance; (Middle) power-dependent 3PAF growth rates under chirped and unchirped/baseline pSLAM illuminations that reveal nonlinear phototoxicity; (Bottom) lower WHF-revealed phototoxicity but early bubble formation (arrowhead) at a higher pSLAM power. **d** (Top) Tri-period eSLAM imaging of chicken breast that reveals linear phototoxicity at a low irradiance; (Bottom) Similar tri-period imaging that reveals the emergence of nonlinear (photoionization-accelerated) phototoxicity and existence of heating-accelerated phototoxicity at a higher irradiance. **e** FLIM imaging of a chicken breast sample containing eSLAM imaging-induced fluorescent compounds at a high irradiance, with color bar corresponding to lifetime in ns. **f** (Left) Illustration of spatiotemporal bin-3 illumination in comparison to baseline/bin-1 illumination; (Right) comparison of 2PAF/3PAF growth rate per frame/pulse and SHG/THG intensity per pulse under bin-1/3/10 pSLAM illumination. **g** Determination of water optical breakdown threshold (arrowhead) by pSLAM baseline ‘imaging’ of 10-mM NADH solution. **h** Simplified view of phototoxicity and related thresholds versus power/irradiance free of cumulative multi-pulse effect.

This result is surprising because muscle samples such as chicken breast are not known as pigmented tissue, in contrast to mouse retina wherein the observed “luminescent flash” can be attributed to linear absorption/phototoxicity^**31**^. It should be noted that two-photon “photo-enhancement” of rabbit red blood cells occurred more readily with longer pulses, implying a photon order of less than 2 (Supplementary Table 5). Moreover, the observed linear phototoxicity via WHF assay echoes that observed from mouse brain slices via the calcium microdomain hyperactivity in cortical astrocytes^**32**^, which was attributed to the heating toxicity that can be detected offline after *in vivo* mouse brain imaging^**33**^.

### Hyper-fluorescence originates from unrelaxed triplet rather than heating

To assess the plausible role of heating in WHF, we lowered the line rate of fast scanning direction (*x*) in the baseline illumination at one pulse per pixel (diffraction-limited resolution of ∼0.4 μm) to bin 3 pulses spatially (Extended Data Table 2, spatial bin). Thus, one frame of the resulting bin-3 illumination (3 pulses/pixel) took the same time and light dose as three frames of the baseline/bin-1 illumination (1 pulse/pixel) (Fig. 3b, top; Extended Data Fig. 3, right). The constant laser repetition rate (0.83 MHz) separated successive pulses apart by 1.2 μs, a period much longer than the thermal relaxation time of water (0.06-0.14 μs) ^**7**,**34**^. Thus, any heating effect in these illuminations would be dominated by single-pulse heating with no interference between successive pulses, which would form the starting point to reveal plausible cumulative multi-pulse heating by increasing the repetition rate^**7**^. In other words, WHF growth rates of the bin-3 illumination would approximate those of the baseline illumination.

Surprisingly, observed 2PAF (3PAF) growth rate in the bin-3 illumination is 4.8-time (3.4-time) of that in the baseline illumination (Fig. 3b, bottom; Extended Data Fig. 3, right), indicating a non-heating cumulative multi-pulse effect. Consistently, the WHF power threshold in the bin-3 illumination is systematically lower than that of the baseline illumination. It is thus unlikely that the observed WHF originates from direct heating^**32,33,35**^. In the context of lowered phototoxicity at increased scanning speed^**17-19**^, the simplest alternative interpretation of this effect is the unrelaxed triplet after the linear absorption of specific intrinsic NIR photosensitizers, with a μs -scale lifetime prone to the excitation state absorption of successive pulses and subsequent high-order triplet-mediated ROS production^**16**^ (Fig. 1d). In contrast to the bin-3 illumination, the baseline illumination at 1 pulse/pixel avoids this ROS production by a faster scanning that spatially separates the individual pixels (excited photosensitizers in one cycle apart by ∼1 diffraction-limited resolution) without the cumulative multi-pulse effect (Fig. 3b, top), and thus relaxes the first-order triplet (Fig. 1d). Without this scanning in a laser tweezer largely free of temperature rise and heating^**36**^, cw-NIR excitation with unrelaxed triplet^**16**^ has induced hyper-fluorescence^**37**^ and ROS-mediated phototoxicity^**20**^ to a trapped cell after a prolonged exposure (∼10 min).

### Photoionization accelerates hyper-fluorescence at high irradiance

To evaluate the power-dependence of WHF growth rates, we employed the baseline and chirped bin-1 pSLAM illuminations (free of the cumulative multi-pulse effect), except for larger powers up to ∼2-time of the corresponding WHF power thresholds. Interestingly, both illuminations led to a nonlinear WHF phototoxicity with an apparent photon order within 3-6 (Extended Data Fig. 4).

**Fig. 4.**
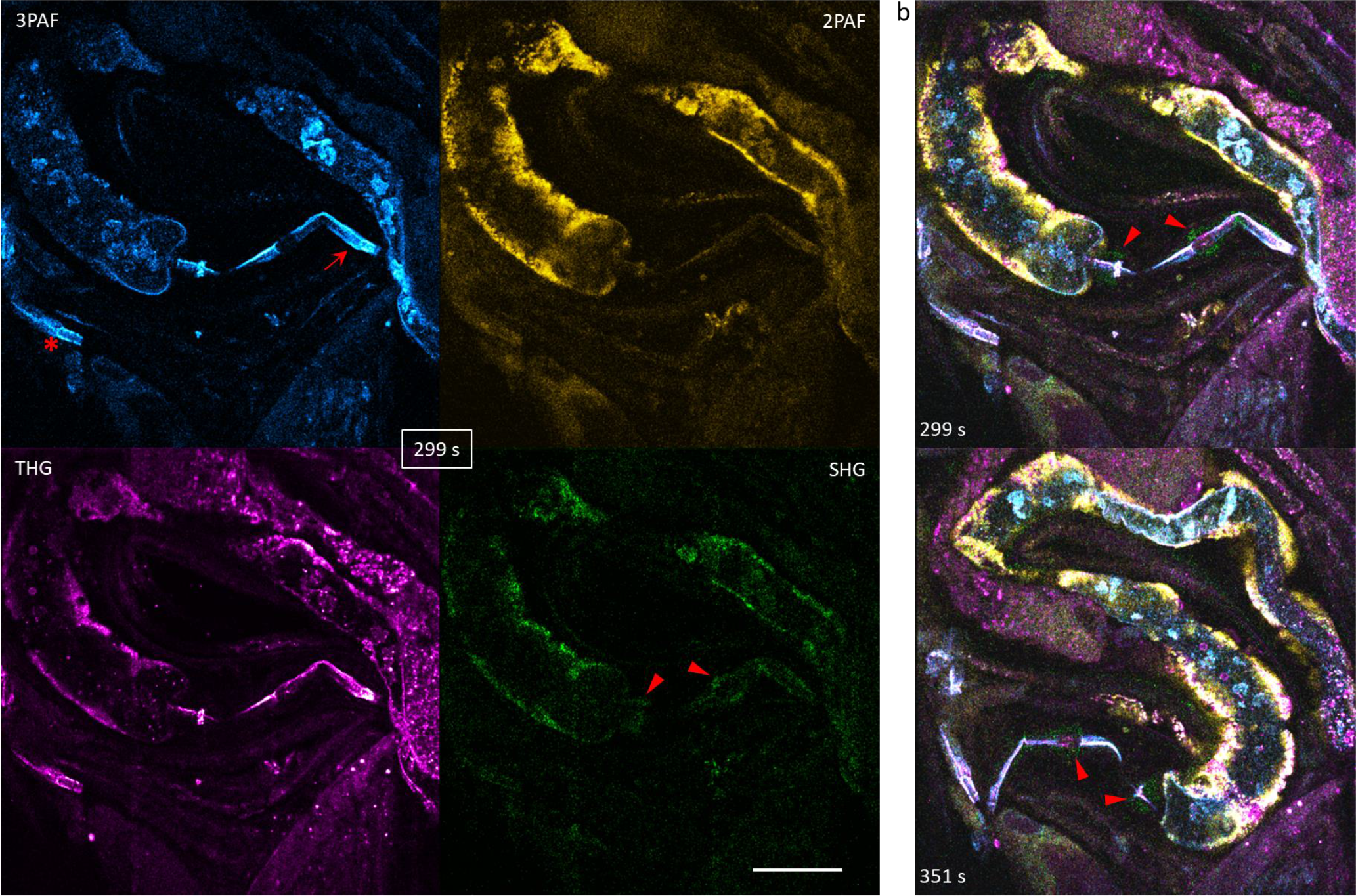
Gentle eSLAM imaging of unlabeled moving *C. elegans* (14.4 mW on sample). **a** Instant single-frame images of one worm with 3PAF/cyan-visible pharynx (arrow) and a pair of SHG/green-visible bulbs (arrowheads); similar pharynx structure (star) indicate the presence of multiple worms in the same field of view. **b** related time-lapse imaging of the worm (arrowheads) with 0.33 s exposure per frame and 1.37 s between successive frames; related video reveals THG/magenta-visible embryos and uterus, 3PAF/cyan-visible proximal and distal gonads with internal germ cells or oocytes, 3PAF/THG-visible intestine, and 2PAF/yellow-visible body wall muscle (Supplementary Video 5). Scale bar: 50 μm.

To identify the factor that accelerates the otherwise linear phototoxicity of WHF toward the nonlinear phototoxicity, we examine the dynamics of high phototoxicity well above the WHF power threshold (Fig. 3c, middle). At the highest power of the baseline illumination, long-lasting bubbles^**10**^ formed near the 80^th^ frame (Fig. 3c, top), whereas a lower power did not form the bubbles after 120 frames (Extended Data Fig. 5, left). In contrast, at the highest power of the chirped illumination, the bubble formation occurred at an earlier (∼15^th^) frame, but the growth rate from either 3PAF or 2PAF was at least 50% lower (Fig. 3c, bottom; Extended Data Fig. 5, right). Given a common temperature threshold for bubble formation (∼130 ºC)^**7**,**10**^, the latter should sustain a higher global temperature^**33**^ than the former throughout the imaging. Thus, WHF is mainly accelerated by single-pulse photoionization dictated by irradiance (unit: TW/cm^2^), not the heating dictated by average power (unit: mW)^**31**^. The resulting photoionization-accelerated WHF might overwhelm the otherwise progressively decreased SHG signal (Fig. 3c, bottom) by thermal denaturing^**30**^ (arrow in Fig. 3c, top).

### Heating accelerates hyper-fluorescence at low irradiance

To generalize the results from pSLAM, we employed the eSLAM microscope that replaced the prism-based compressor with a pulse shaper to electronically chirp 1110-nm pulses from 300 fs to 60 fs (Extended Data Table 2)^**29**^. With the same spatially separated pulsed excitation (i.e., one-pulse-per-pixel imaging of the same FOV), the observed WHF power threshold increased from 1.8-mW in pSLAM to 17-mW in eSLAM (Extended Data Fig. 1). However, the corresponding irradiances (i.e., 2.2-nJ pulse energy at 1030-nm in pSLAM versus 3.4 nJ at 1110-nm in eSLAM) are comparable, indicating the single-pulse nature of phototoxic WHF (free of cumulative multi-pulse effect). This single-pulse effect also dominates the cumulative multi-pulse effect in bubble formation or heating toxicity (Extended Data Fig. 5, right). Using a power 10% above the WHF power threshold (Extended Data Table 2, eSLAM), we conducted a tri-period imaging with switchable pulse chirping and observed the same linear WHF growth rate in the first and third periods, which was sustained by the second period even though the corresponding phototoxicity could not be detected by chirped pulses (Fig. 3d, top). This independently confirms the linear absorption nature of WHF observed in pSLAM (Fig. 3a).

To examine the plausible role of heating, we used a power ∼20% above the WHF power threshold and observed a much larger WHF growth rate from the first period than that from the second period, which indicated the emergence of photoionization-based acceleration (Fig. 3d, bottom). Interestingly, the WHF growth rate from the third period surpassed that from the first period considerably (Fig. 3d, bottom). If linearly absorbed light energy end ups more as heat than the photoionization-accelerated WHF during the second period in comparison to the first period, we attribute the higher WHF growth rate of the third period than that of the first period to a higher global temperature^**33**^. This implies that the global heating, which is superimposed on the single-pulse heating in eSLAM (with 0.2 μs pulse separation larger than the thermal relaxation time of 0.06-0.14 μs), accelerates WHF under a relatively low irradiance (Fig. 1d). This acceleration may be caused by the thermal inactivation of intrinsic ROS scavengers in addition to an increased ROS production^**23**^. As an implication, it allows detection of linear phototoxicity by a chirping-pulse experiment (Fig. 3a, bottom; Fig. 3d, top) but not a conventional power-dependence experiment (Fig. 3c, middle; Extended Data Fig. 4).

### Hyper-fluorescence lifetime generalizes model applicability

To measure the lifetime of WHF, we invoked the fluorescence lifetime imaging microscopy (FLIM) capability of the eSLAM microscope^**29**^. The WHF was first induced in one FOV (∼15 μm imaging depth) by 200 scans at a rather high power to overwhelm the intrinsic fluorescence signal, and then FLIM imaging was conducted on the same FOV using a low power that did not further increase the WHF. The lifetime of homogeneous WHF (∼1.5 ns) is longer than that of heterogenous WHF (∼0.6 ns; Fig. 3e, arrowheads), which are comparable with those observed from cultured hamster kidney cells and *ex vivo* mouse kidney (Fig. 2a,b; Extended Data Fig. 6a-6c). Similar lifetimes (∼1 ns) were obtained from the NIR-induced “white flashes” of Chinese hamster cells^**38**^ and from new fluorescent compounds in NIR-ablated muscle and albumin samples^**35**^, suggesting that similar fluorescent compounds produce the widely reported WHF in diverse cell/tissue samples (Supplementary Table 5).

The homogeneous WHF was also observed in a mouse brain slice via 3PAF but not 2PAF across the FOV, whereas the latter could reveal this WHF in a central region of FOV with the highest irradiance (Extended Data Fig. 6d; Supplementary Video 4). From this result, we believe the photobleaching of strong intrinsic fluorescence often obscures the homogeneous WHF, which would otherwise be widely observable in live samples. The intrinsically weak 2PAF/3PAF signals of chicken breast offer a rather clean background at the beginning of time-lapse imaging for sensitive detection of the homogeneous WHF and accurate quantification of the phototoxicity.

### Cumulative multi-pulse effect lowers imaging performance

In contrast to the spatial bin-3 illumination that binned 3 pulses into one spot spatially (Fig. 3b, top), the more popular binning affordable by the flexible galvo-galvo scanning and tunable repetition-rate laser of pSLAM was to bin multiple pulses into one spot spatiotemporally at the same fast-axis scanning speed of the baseline illumination^**28**^ (Fig. 2c,d; Fig. 3f, left). To evaluate the benefit/cost of this spatiotemporal binning, we performed imaging on different FOVs at bin-1 (baseline), bin-3, and bin-10 conditions with different powers (repetition rates) but the same pulse energy (Extended Data Table 2). Unlike the spatial binning case (Fig. 3b, bottom), the WHF growth rate per pulse decreases with the bin number (Fig. 3f, upper right) possibly due to saturated absorption of the photosensitizers. However, this benefit is countered by the cost of lowered nonlinear signals (Fig. 3f, lower right), which may be attributed to unrelaxed photoionization with up to 300 ns lifetime of solvated electrons^**10**^. The corresponding cumulative multi-pulse effect produces similar nonlinear phototoxicity (Extended Data Fig. 4). However, it lowers the WHF irradiance threshold due to a higher WHF growth rate per frame (Fig. 3f, upper right). Thus, it is more beneficial to increase the throughput of the pSLAM baseline imaging by the bin-1 (no-bin) illumination of eSLAM, free of the cumulative multi-pulse effect that likely impaired embryo development^**39**^. This eSLAM illumination relaxes the linear-absorption-mediated triplet (Fig. 1b) by combining a higher repetition-rate excitation with a proportionally faster scanning, and thus attains the same one-pulse-per-pixel imaging across the same FOV (Extended Data Table 2).

To estimate the optical breakdown threshold of water, we conducted the pSLAM baseline illumination on a 10-mM solution of reduced nicotinamide adenine dinucleotide (NADH) at different pulse energies and identified a threshold of 5.3-nJ (Fig. 3g) corresponding to 9.3 TW/cm^2^ at 1030-nm (near identical threshold was attained at 1110-nm). This threshold approximates the theoretical value of 3.9-nJ or ∼7 TW/cm^2^ (1064-nm, 100-fs, NA 1.3)^**10**^. Thus, gentle imaging free of unrelaxed triplet and subsequent ROS production (detectable WHF in Extended Data Fig. 1) can be conducted at a surprisingly large ratio (∼50%) of water breakdown threshold (normalized irradiance in Extended Data Table 3). The irradiance/power-dependent phototoxicity becomes clear without the interfering cumulative multi-pulse effect (Fig. 3h).

### Triplet-relaxation and fast scanning enables high-performance imaging

To highlight the ‘gentle’ eSLAM imaging operated near one-half of water breakdown threshold (Extended Data Table 3), we investigated wide-type unlabeled *C. elegans* in standard culture (OP50). All signals were epi-detected and displayed at 0.73 Hz frame rate (1024×1024 -pixel frame) without image denoising/reconstruction. We observed the free motion of one worm with 0.2-μs pixel dwell time and identified its moving 3PAF-visible pharynx and related pair of SHG-visible bulbs (Fig. 4; Supplementary Video 5). Similar results, including the SHG-visible bulbs, were obtained in less adaptive transmission configuration with 5-μs pixel dwell time that necessitated animal immobilization^**40**^. More importantly, it is unprecedented to observe strong back-reflected SHG signal from transparent samples such as *C. elegans*, which is worth further fundamental study. This demonstration highlights the potential of eSLAM to translate label-free nonlinear optical imaging (Supplementary Table 3).

## Discussion

The coexistence of linear and nonlinear NIR phototoxicity observed in our WHF assay (at low and high powers, respectively) echoes the similar observation in a previous study at ∼30-fold lower irradiance^**41**^ (Extended Data Table 3) and reconciles the existence of either linear or nonlinear NIR phototoxicity in various WHF-like assays (Supplementary Table 5) and bioassays (Supplementary Table 6). Apparently nonlinear phototoxicity originates from the linear NIR absorption^**32**,**41**^ of ubiquitous intrinsic NIR photosensitizers^**20-23**^, just like that from the excessive (high-power) linear absorption of intrinsic UV-visible photosensitizers^**32**^. Thus, the apparent photon order of ≥2 in the observed phototoxicity (Supplementary Tables 5,6) does not imply a multiphoton absorption, as would be commonly assumed (Fig. 1b). In other words, the concept of photon order is useful to characterize the nonlinearity of molecular absorption and harmonic generation but not a phenomenological phototoxicity. Without this “evidence” that has strongly supported the nonlinear-absorption-mediated phototoxcity, other evidence can be reinterpreted to be compatible with our hypothesized mechanism of linear-absorption-mediated nonlinear phototoxicity (Supplementary Table 4). To avoid the relevant accelerated ROS production in (clinical) nonlinear optical imaging (Supplementary Table 3), it is mandatory to relax the triplet state by increasing the fast-axis scanning speed (or pulses per diffraction-limited-resolution PPD; see Supplementary Table 2), just like in linear optical imaging^**17-19**^. The increased scanning speed allows real-time sensitive monitoring of phototoxicity via WHF^**24**^ and unambiguous identification of a non-fluence (power, irradiance, intensity, etc.) threshold with minimal doses (from first frames in Extended Data Fig. 1). Therefore, our study may tip the balance from the popular view of pulsed-NIR phototoxicity based on a fluence threshold (Supplementary Table 1) toward the contradictory view based on an irradiance threshold (Fig. 3h).

One major confusion on observed NIR phototoxicity (Supplementary Tables 1,4) arise from the existence of different relaxation time scales of >1 μs (photochemistry), ∼0.3 μs (photoionization), and ∼ 0.1 μs (heating) to produce different extents of cumulative multi-pulse effect (Figs. 3b,3f). This effect is prevalent in the typical illumination (galvo-galvo scanning of ∼80 MHz pulses) of nonlinear optical imaging (Supplementary Table 2) and may have obscured the hypothesized mechanism. By resonant-galvo scanning of ∼5 MHz pulses to remove this effect (so that subsequent pulses address well-resolved pixels^**16**^ ∼1 diffraction-limit resolution apart, i.e., triplet-relaxation in eSLAM), gentle imaging can be conducted at ∼50% of water optical breakdown before the linear/nonlinear phototoxicity onset of single-pulse heating/photoionization (Figs. 3d,3h). Without this relaxation, laser surgery can occur at only 8.6% of water optical breakdown^**42**^ (Extended Data Table 3). Also, accelerated phototoxicity by cumulative multi-pulse heating^**8**,**38**^ may induce linear-absorption-mediated nonlinear phototoxicity^**32**^ at ≤3% of water optical breakdown (largely free of single-pulse photoionization), resembling the single-pulse heating in the picosecond excitation of coherent anti-Stokes Raman scattering microscopy with this relaxation^**43**^ (Extended Data Table 3). This critical role of single-pulse heating (Supplementary Table 1) is further validated by the low phototoxicity onset below 10% of water optical breakdown in pigmented specimens of skin^**7**^ and retina^**31**^ (Extended Data Table 3), and the absence of reported WHF in low-power UV-visible confocal fluorescence microscopy. However, our hypothesized mechanism attributes the corresponding linear phototoxicity more to unrelaxed triplet and subsequent heating-accelerated ROS production (Fig. 1d) than direct heating^**7,31,32,35**^.

We have tested the eSLAM irradiance threshold from chicken breast model (Extended Data Table 3) in diverse cultured cells and *ex vivo* or *in vivo* mouse tissue specimens. An irradiance 10% more than this threshold consistently generates WHF in time-lapse imaging (Figs. 2a,2b) whereas an irradiance 10% less than this threshold avoids the WHF completely. This test not only validates the assertion of a non-fluence threshold below which dose becomes irrelevant^**14**^, but also the ability of WHF (or chicken breast) as a real-time inline indicator^**24**^ (or a ‘natural’ tissue-mimicking phantom^**44**^) for phototoxicity. The intrinsic WHF avoids the limitations of the photobleaching of specific extrinsic fluorophores^**2**,**3**^ to indicate subtle/early phototoxicity and restricts more severe phototoxicity of damage to nucleus^**8**,**24**^ and morphology,^**38,41**^. Thus, short (sub-80-fs) pulse^**31**,**45**^ is preferred over longer (picosecond) pulses^**41**,**43**^ to not only increase nonlinear signal generation near linear phototoxicity onset^**32**^ but also detect this onset more sensitively than 300-fs pulses (Fig. 3d, top) within a narrow window of linear phototoxicity (Fig. 3h). As to the pulse repetition rate for triplet-relaxation, the choice depends on the trade-off between imaging depth^**45**^ (5-MHz of eSLAM preferred) and speed^**33**^ (20-MHz fast version of eSLAM preferred; see Extended Data Table 2), both of which are subjective to the constraint of global heating^**33**^. The optimal repetition rate to balance this tradeoff likely lies within the 5-20 MHz range. Either way, our hypothesized mechanism offers a universal technique to mitigate the NIR phototoxicity (Fig. 1d) absent from the existing mechanism (Fig. 1b). Further improvements may enable a more complete triplet-relaxation without spatial under-sampling^**46**^ and a higher throughput via volumetric imaging^**47**^, while neutralizing heat/ROS generation may mitigate phototoxicity *in vitro*^**20**,**23**^.

For pigmented samples, 3PAF imaging of NDAH at 1110-nm (eSLAM) may outperform conventional 2PAF imaging of NDAH at 750-780 nm^**7**,**31**^ due to low linear absorption of melanin (single-pulse heating) at longer wavelengths^**48**^. For general (non-pigmented) biological samples, more studies on biochemistry are needed to determine whether there exists an optimal wavelength across NIR (700-1300 nm) for gentle nonlinear optical imaging (Supplementary Table 3) and an action spectrum of phototoxicity^**20**,**21**^. The emergence of WHF reveals the otherwise invisible myofibrils^**49**^ in chicken breast (Fig. 2d), suggesting an origin of fluorescent Schiff base in lipofuscin-like products from the peroxidation of polysaturated lipids that crosslinks proteins^**24,25,50**^. It would be interesting to generalize the WHF as a form of oxidation-induced fluorescence from light exposure, paralleling that from heating/cooking^**35**^, storage-induced meat deterioration^**49**^ or natural aging^**50**^, and chemical induction^**51**^. However, the dependence of NIR phototoxicity on wavelength may rely more on the absorption properties of photosensitizers (initiator) and plausible heat-generating pigments or chromophores (accessory) than those of the resulting species emitting WHF (end-product) (Fig. 1d). In summary, with rapidly scanned sub-80-fs excitation to fully relax the triplet state, the unfulfilled potential^**3**^ of laser-scanning nonlinear optical microscopy^**6**^ in gentle imaging^**52-54**^ may be rationally recovered after its early demonstration in 1990.

**Extended Data Table 1.** Comparison of heterogeneous and homogeneous WHF in SLAM-based imaging.

|  | Heterogeneous WHF | Homogeneous WHF |
| --- | --- | --- |
| Prevalence | Inevitably present in cell culture and live tissue at sufficiently high power/irradiance or dosage | Observable in *ex vivo* chicken breast and mouse brain but not cell culture and *ex vivo* mouse kidney |
| Initial photodamage site | Random point(s) within the field-of-view | Center of field-of-view with the highest irradiance |
| Time-lapse morphology | Increase of both intensity and spatial scope in a point-spreading fashion | Uniform elevation across a large area of illumination field |
| Nature of phototoxicity | Dependent on light dosage like bubble formation by ultrashort pulses | Dependent on threshold power/irradiance unlike the bubble formation by ultrashort pulses |
| Time-lapse phototoxicity | Not observable in the first frame but with nonlinear increase over frame/time (Extended Data Fig. 6b,c) | Observable in the first frame with linear increase over frame/time |
| Stage of phototoxicity | Late stage after onset of homogeneous WHF (chicken breast as example) | Early stage during photobleaching-to-WHF transition (chicken breast and mouse brain as examples) |
| Lifetime | Short (~0.6 ns) | Long (>1 ns) |
| Quantitative analysis and modeling of phototoxicity | Unsuitable due to random and point-spreading morphology, dependence on light dosage, and nonlinear increase at a late stage of phototoxicity | Suitable due to homogeneous morphology, dependence on threshold power/irradiance, and linear increase at an early stage of phototoxicity |

**Extended Data Table 2.** Illuminations of pSLAM and eSLAM on chicken breast.

| Microscope (optical scanner, control software) | pSLAM (galvo-galvo, ScanImage) |  |  |  | eSLAM (resonant-galvo, LabVIEW) |
| --- | --- | --- | --- | --- | --- |
|  | Baseline/unchirped | Chirped | Spatial bin-3 | Spatiotemporal bin-10 (or bin-3) |  |
| Central wavelength | 1030 nm | 1030 nm | 1030 nm | 1030 nm | 1110 nm |
| Pulse width on sample (FWHM) | 60 fs | 300 fs | 60 fs | 60 fs | 60 fs unchirped (or 300 fs chirped) |
| Bin: pulses/pixel/frame | 1 | 1 | 3 | 10/3 | 1 |
| Illumination *P* on sample | 2.0 mW | 4.0 mW | 2.0 mW | 20/6.0 mW | 18.7 mW |
| Pulse repetition rate | 0.83 MHz | 0.83 MHz | 0.83 MHz | 8.3/2.5 MHz | 5 MHz |
| Fast scan line rate | 340 Hz | 340 Hz | 113 Hz | 340 Hz | 1592 Hz |
| Pixel dwelling time | 1.4 μs | 1.4 μs | 4.2 μs | 1.4 μs | 0.2 μs |
| Exposure/acquisition time per frame | 1.5/1.5 s | 1.5/1.5 s | 4.5/4.5 s | 1.5/1.5 s | 0.33/1.37 s |
| Relevant figure(s) in main text | Figs. 3a,3b,3f | Fig. 3a | Fig. 3b | Fig. 3f | Fig. 3d, top |

Note on common features: imaging depth ~10 μm; frame 1024 pixel × 1024 pixel; field-of-view 300 μm × 300 μm.

**Extended Data Table 3.**
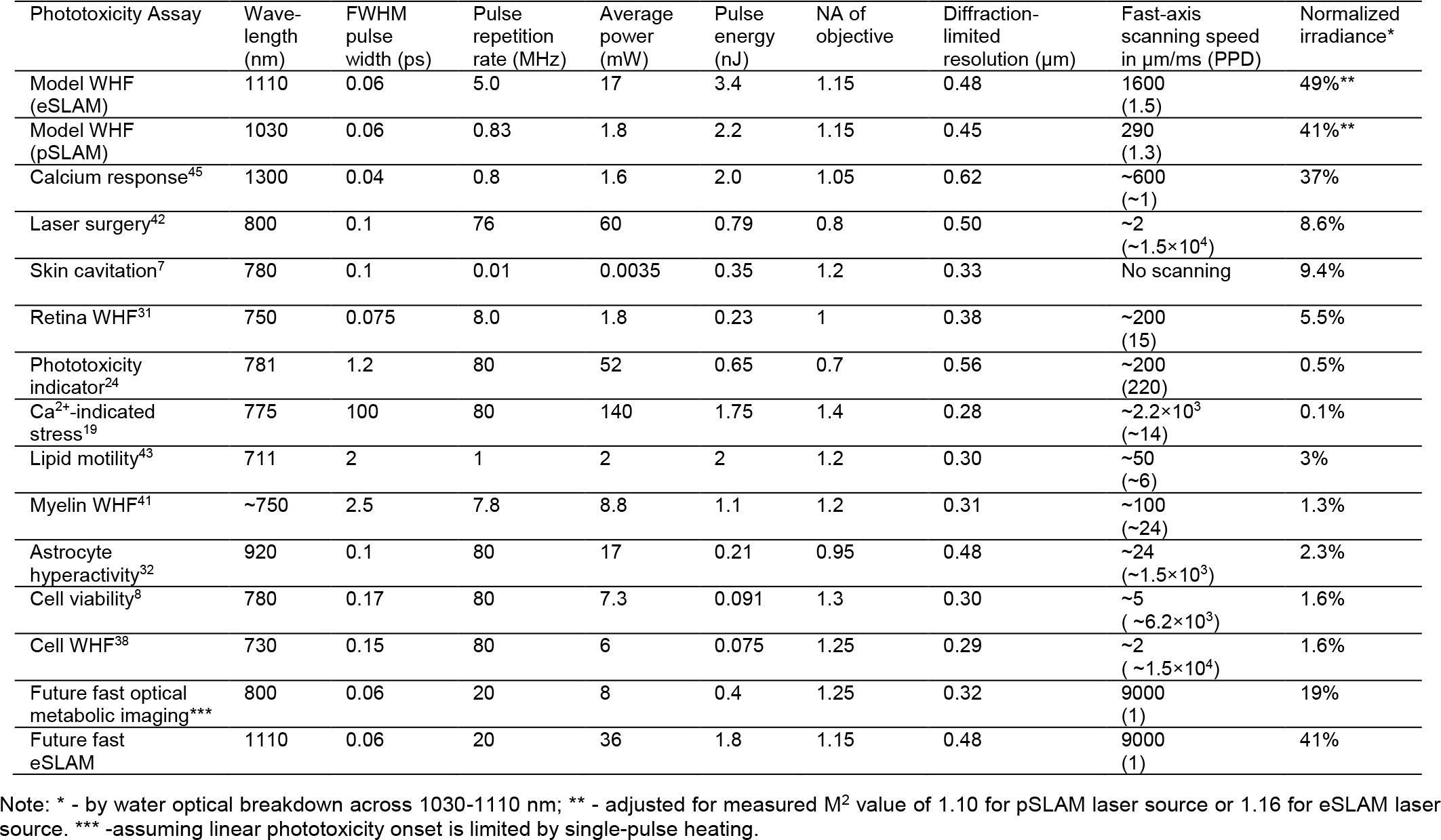
Normalized threshold irradiance of NIR phototoxicity versus number of pulses per diffraction-limited-resolution.

| Phototoxicity Assay | Wave-length (nm) | FWHM pulse width (ps) | Pulse repetition rate (MHz) | Average power (mW) | Pulse energy (nJ) | NA of objective | Diffraction-limited resolution (μm) | Fast-axis scanning speed in μm/ms (PPD) | Normalized irradiance* |
| --- | --- | --- | --- | --- | --- | --- | --- | --- | --- |
| Model WHF (eSLAM) | 1110 | 0.06 | 5.0 | 17 | 3.4 | 1.15 | 0.48 | 1600 (1.5) | 49%** |
| Model WHF (pSLAM) | 1030 | 0.06 | 0.83 | 1.8 | 2.2 | 1.15 | 0.45 | 290 (1.3) | 41%** |
| Calcium response<sup>45</sup> | 1300 | 0.04 | 0.8 | 1.6 | 2.0 | 1.05 | 0.62 | ~600 (~1) | 37% |
| Laser surgery<sup>42</sup> | 800 | 0.1 | 76 | 60 | 0.79 | 0.8 | 0.50 | ~2 (~1.5×10<sup>4</sup>) | 8.6% |
| Skin cavitation<sup>7</sup> | 780 | 0.1 | 0.01 | 0.0035 | 0.35 | 1.2 | 0.33 | No scanning | 9.4% |
| Retina WHF<sup>31</sup> | 750 | 0.075 | 8.0 | 1.8 | 0.23 | 1 | 0.38 | ~200 (15) | 5.5% |
| Phototoxicity indicator<sup>24</sup> | 781 | 1.2 | 80 | 52 | 0.65 | 0.7 | 0.56 | ~200 (220) | 0.5% |
| Ca<sup>2+</sup>-indicated stress<sup>19</sup> | 775 | 100 | 80 | 140 | 1.75 | 1.4 | 0.28 | ~2.2×10<sup>3</sup> (~14) | 0.1% |
| Lipid motility<sup>43</sup> | 711 | 2 | 1 | 2 | 2 | 1.2 | 0.30 | ~50 (~6) | 3% |
| Myelin WHF<sup>41</sup> | ~750 | 2.5 | 7.8 | 8.8 | 1.1 | 1.2 | 0.31 | ~100 (~24) | 1.3% |
| Astrocyte hyperactivity<sup>32</sup> | 920 | 0.1 | 80 | 17 | 0.21 | 0.95 | 0.48 | ~24 (~1.5×10<sup>3</sup>) | 2.3% |
| Cell viability<sup>8</sup> | 780 | 0.17 | 80 | 7.3 | 0.091 | 1.3 | 0.30 | ~5 (~6.2×10<sup>3</sup>) | 1.6% |
| Cell WHF<sup>38</sup> | 730 | 0.15 | 80 | 6 | 0.075 | 1.25 | 0.29 | ~2 (~1.5×10<sup>4</sup>) | 1.6% |
| Future fast optical metabolic imaging*** | 800 | 0.06 | 20 | 8 | 0.4 | 1.25 | 0.32 | 9000 (1) | 19% |
| Future fast eSLAM | 1110 | 0.06 | 20 | 36 | 1.8 | 1.15 | 0.48 | 9000 (1) | 41% |

Note: \* - by water optical breakdown across 1030-1110 nm; \*\* - adjusted for measured M<sup>2</sup> value of 1.10 for pSLAM laser source or 1.16 for eSLAM laser source. \*\*\* - assuming linear phototoxicity onset is limited by single-pulse heating.

**Extended Data Fig. 1.**
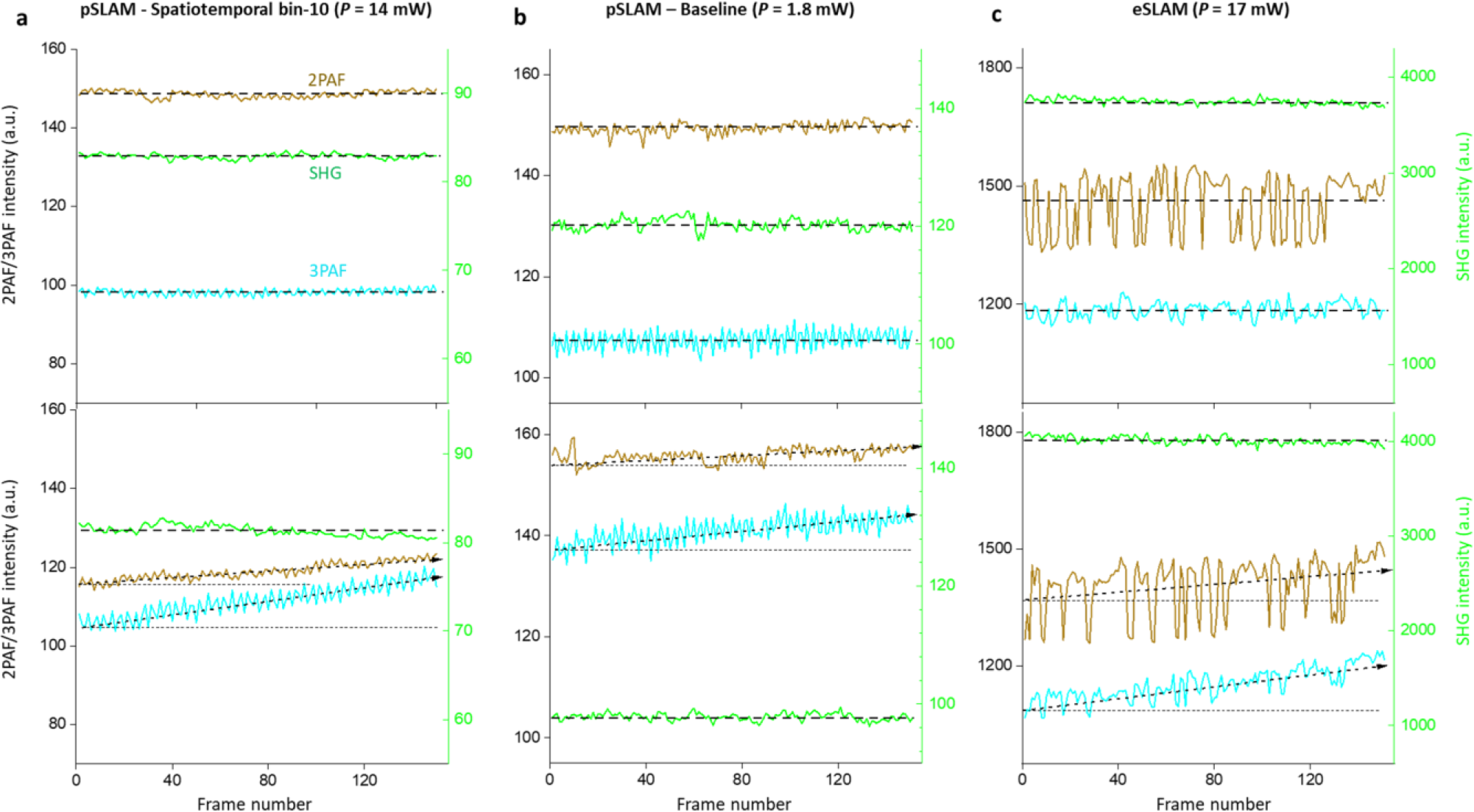
Power threshold to generate WHF in chicken breast model by pSLAM (left two panels) or eSLAM (right panel). At the power threshold, spatially integrated 2PAF and 3PAF (i.e., WHF) signals may not (upper panel) or may (lower panel) increase with frame number during time-lapse imaging, using spatially integrated SHG signal as reference. Details of three illumination conditions are shown in Extended Data Table 2 except for lower powers.

**Extended Data Fig. 2.**
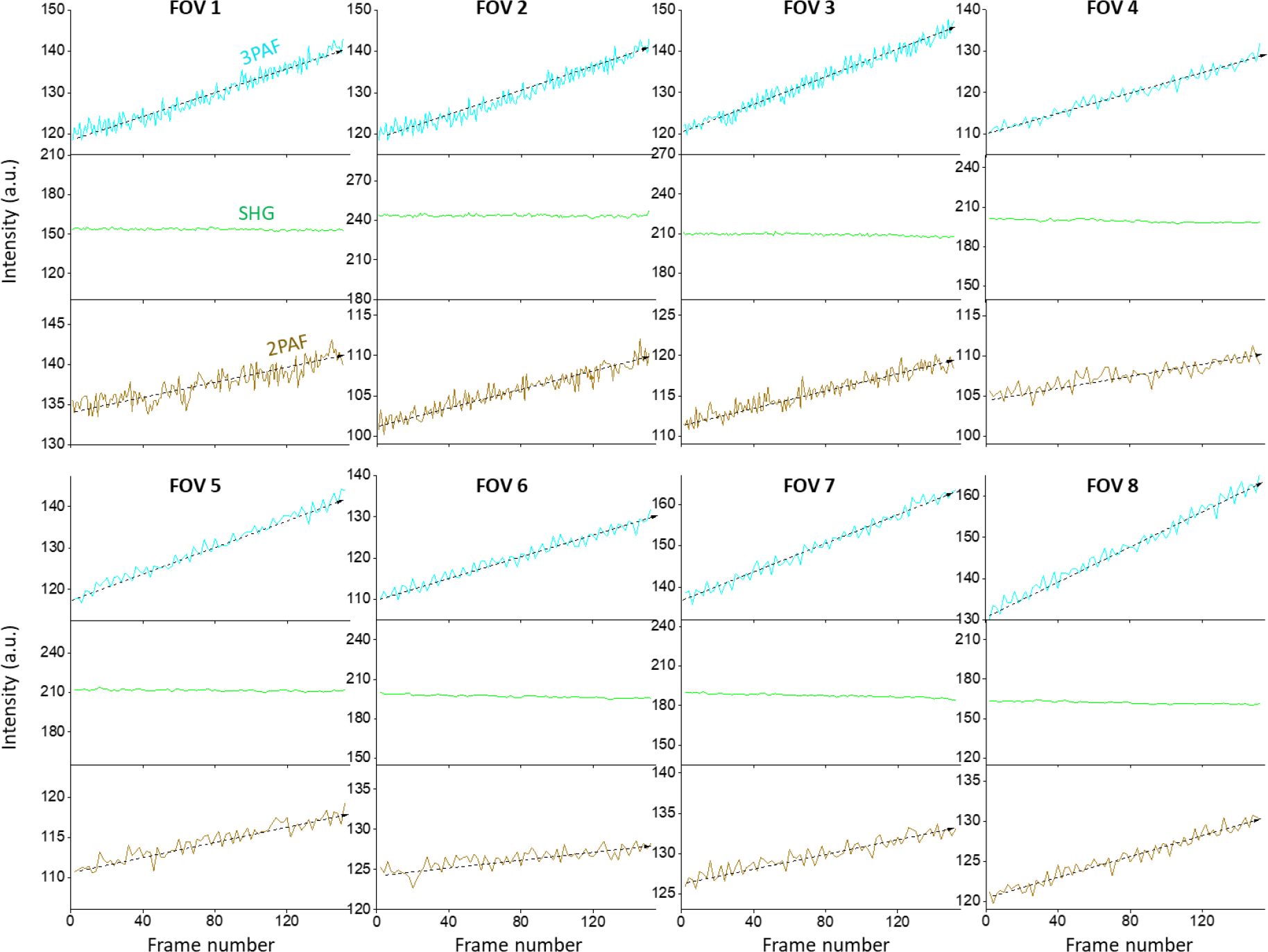
Reproducible 3PAF and 2PAF growth rates under the baseline pSLAM illumination across different FOVs in chicken breast. Spatially integrated 2PAF, 3PAF, and SHG signals during time-lapse imaging are plotted in the same intensity scales despite their difference in absolute intensity across FOVs.

**Extended Data Fig. 3.**
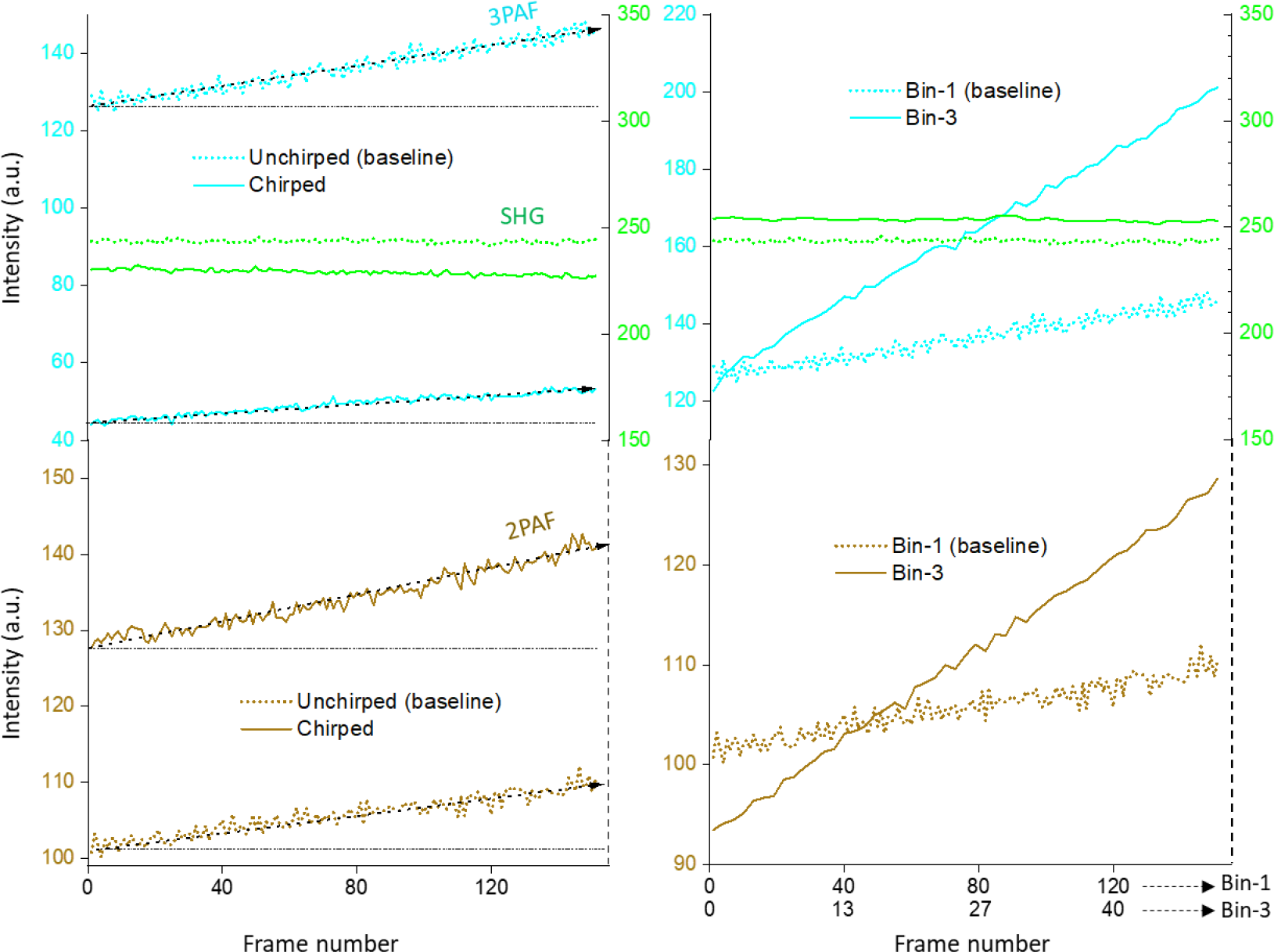
Effect of pSLAM pulse chirping (left) or pulse spatial binning (right) on WHF growth rates. In comparison to unchirped/baseline illumination (60 fs, 2.0 mW), chirped illumination (300 fs, 4.0 mW) attains a higher 2PAF growth rate but a lower 3PAF growth rate. In comparison to bin-1 illumination (baseline), bin-3 illumination (60 fs, 2.0 mW) attains much higher 2PAF and 3PAF growth rates (note that signal intensity of the bin-3 illumination integrated over one frame is normalized by the number of binning).

**Extended Data Fig. 4.**
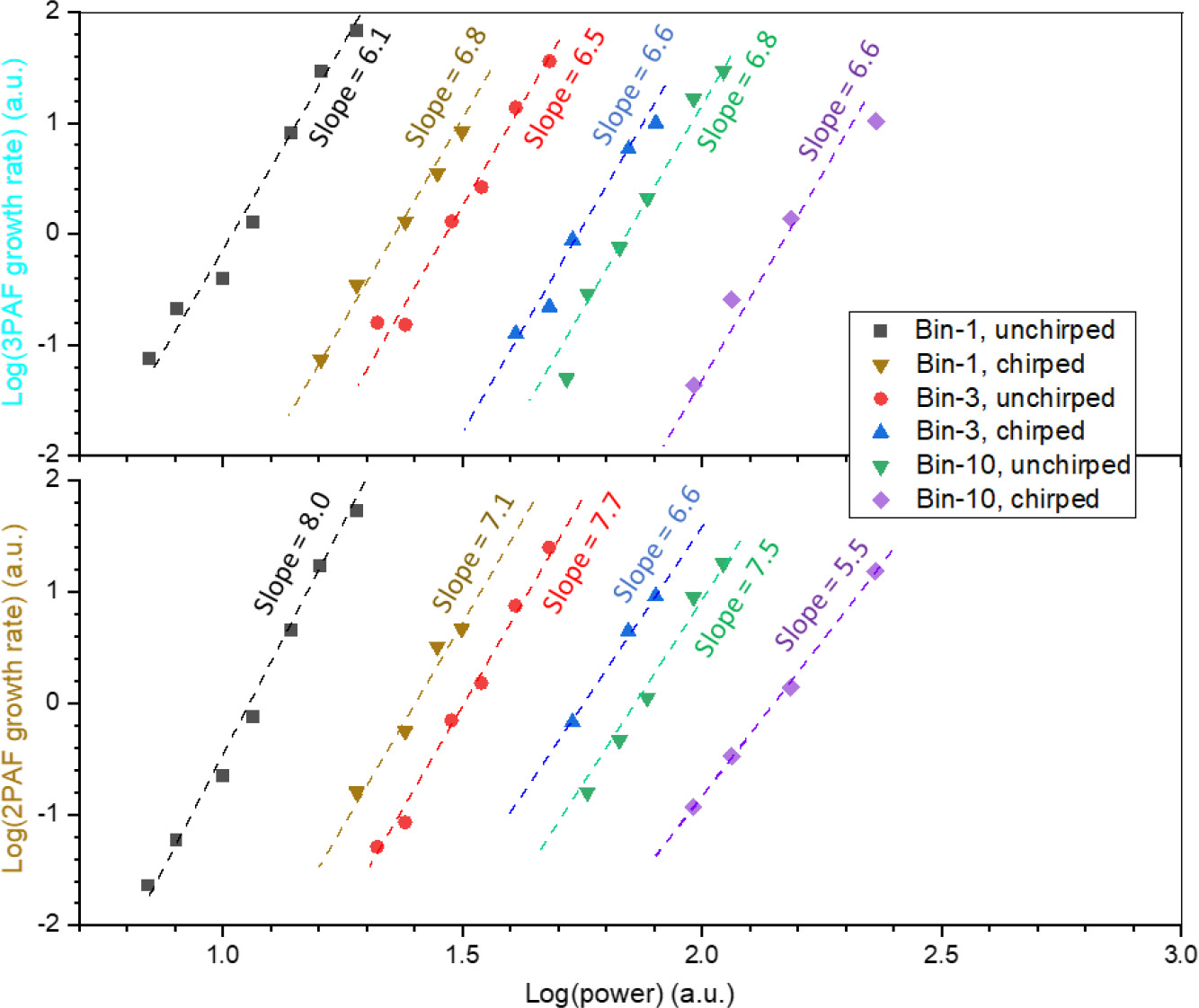
Apparent photon orders of WHF growth rates revealed by 3PAF (top) and 2PAF (bottom) with/without pSLAM pulse chirping or spatiotemporal binning. The apparent photon orders vary across 6.1-6.8 for 3PAF (photon order 3), indicating a nonlinear phototoxicity of order 3.1-3.8. The related apparent photon orders vary across 5.5-8.0 for 2PAF (photon order 2), indicating a nonlinear phototoxicity of order 3.5-6.

**Extended Data Fig. 5.**
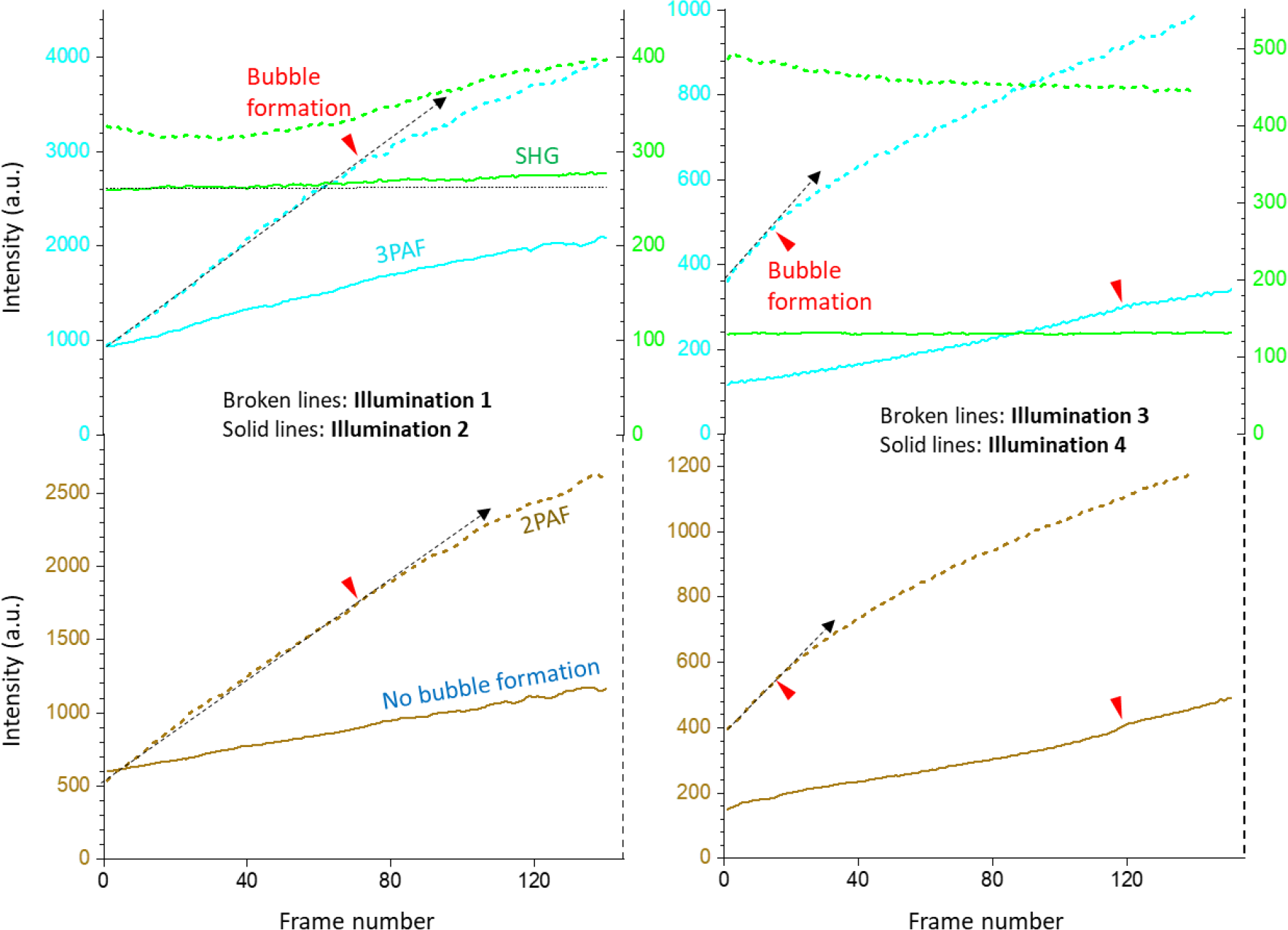
Assessment of bubble formation (arrowheads) under different pSLAM illuminations. (Left) bubble formation at illumination 1 (0.83 MHz 60 fs, 3.0 mW) absent from illumination 2 at a lower power (0.83 MHz 60 fs, 2.6 mW); (Right): bubble formation at illumination 3 (0.83 MHz 300 fs, 5.9 mW) occurs earlies than at illumination 4 (5.0 MHz 300 fs, 20.9 mW) despite the higher power of the latter, indicating the larger role of single-pulse heating than overall thermal load to promote bubble formation. Comparison between Illumination 1 and Illumination 3 (or Illumination 2 and Illumination 4) reveals the larger role of photoionization than heating (relevant to bubble formation) to accelerate 3PAF/2PAF growth rates (i.e., phototoxicity).

**Extended Data Fig. 6.**
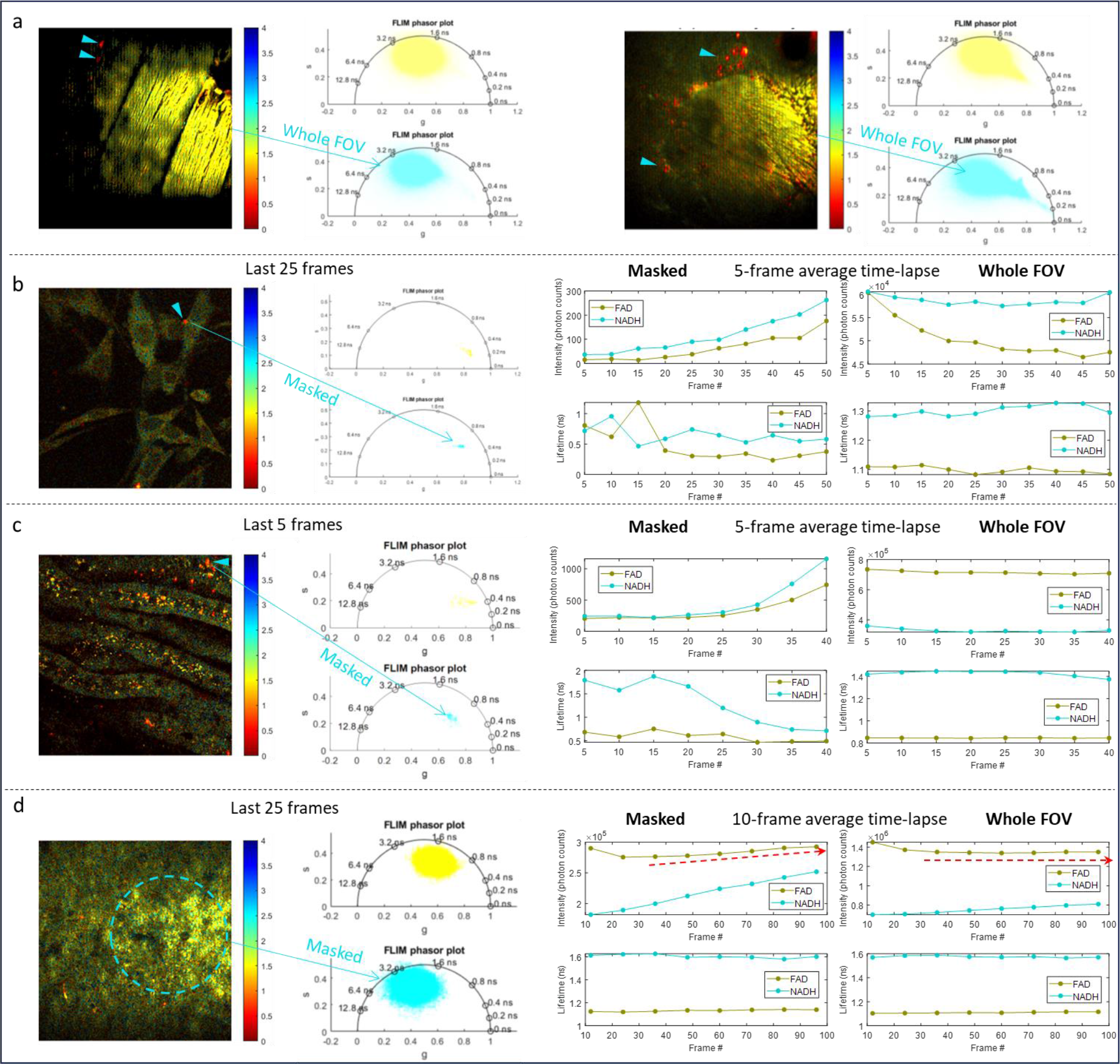
**FLIM of WHF in cultured cells and *ex vivo* tissue** (color bar corresponds to lifetime in ns). **a** Fluorescence lifetime and phasor plot (FAD – yellow; NADH – cyan) observed from two fresh chicken breast samples with homogeneous WHF (yellow center) and punctuated homogeneous WHF (arrowheads). **b** Heterogeneous WHF lifetime (arrowhead) and phasor plot observed from a hamster kidney cell in cell culture (left two panels) at 18 mW; right panels show time-lapse fluorescence intensity and lifetime corresponding to the heterogeneous WHF only (left) and whole FOV (right). **c** Heterogeneous WHF lifetime (arrowhead) and phasor plot observed from *ex vivo* mouse kidney tissue (left two panels) at 18 mW; right panels show time-lapse fluorescence intensity and lifetime corresponding to the heterogeneous WHF only (left) and whole FOV (right). **d** Homogeneous WHF lifetime (broken cycle) and phasor plot observed from *ex vivo* mouse brain slice (left two panels) at 20 mW; right panels show time-lapse fluorescence intensity and lifetime corresponding to the homogeneous WHF only (left) and whole FOV (right).

## Methods

### Imaging instrumentation

Details of our pSLAM microscope and eSLAM microscope have been described in ref. 28 and ref. 29, respectively. Main independent parameters of the pSLAM and eSLAM illuminations are compared in Extended Data Table 2. Average power at the sample plane was measured by a microscope slide power meter (S175C, Thorlabs). The M^2^ value of pSLAM laser source (1.10) or eSLAM laser source (1.16) was measured by a commercial device (M2MS, Thorlabs) to calculate irradiance. All imaging experiments were conducted at room temperature with no additional temperature control of cell/tissue samples.

### Cell culture

Adherent Syrian golden hamster kidney fibroblast cells (BHK-21, clone 13, ATCC #CCL-10) were cultured in disposable BioLite™ 75 cm^2^ vented-cap cell culture treated flasks according to supplier-recommended protocols. They were maintained inside a humidified incubator with 5% CO_2_ and 21% O_2_ conditions at 37°C. A 0.5 -1 ml volume of harvested cells was resuspended in 1.5-1 mL of phenol red-free GibcoTM 1X TrypLE™ Select Enzyme (pH 7.0 – 7.4) cell dissociation reagent (TFS, Cat #12563029) in triplicates. The cells were imaged within 10 min of the resuspension.

### C. elegans

*C. elegans* growing on agar plates seeded with E. coli were obtained from Carolina Biological Supply Company. After additional growth of 2–4 days, a small portion was cut out of the agar plate and placed in a dish (P35G-0-10-C, MatTek) for imaging.

### Animal tissue

Fresh chicken breast was purchased from a local supermarket, cut by a razorblade with a smooth surface, and imaged within 24 hrs. All experiments on rodents were performed in compliance with the Guide for Care and Use of Laboratory Animals of the National Institutes of Health and approved by the Institutional Animal Care and Use Committee at the University of Illinois at Urbana-Champaign (Animal Welfare Assurance #A3118-01). Brains of 4-week Long-Evans rats from an inbred colony (LE/BluGill) were removed and immersed in ice-cold slicing media (93 mM N-Methyl-D-glucamine, 2.5 mM KCl, 1.2 mM NaH2PO4, 30 mM NaHCO3, 20 mM HEPES, 25 mM glucose, 2 mM Thiourea, 3 mM Sodium pyruvate, 10 mM MgSO4, 0.5 mM CalCl2, pH 7.4) bubbled with CO_2_. Coronal slices (400 μm) containing the medial suprachiasmatic nucleus (SCN) were sectioned by a vibratome (Leica VT1000S). Slices were transferred to tissue culture inserts (0.4 μm; Millicell -CM, Millipore) contained within 35mm tissue culture dishes. The dishes were immersed in 1 ml of organotypic media, i.e., DMEM without sodium pyruvate supplemented with 10 mM HEPES, GS21 (1:50, GlobalStem), Penicillin-Streptomycin (1:100, ThermoFisher Scientific) and 1 mM L-glutamine. Cultures were kept at 37 °C i n 5 % CO_2_ and media were exchanged every other day. Brain slices kept in culture for < 1 week were used for imaging. Mice (C57BL/6J, Jackson Laboratory) were used to obtain *ex vivo* kidney samples, which were imaged directly without specific preparation.

## Data availability

The data that support the findings in this study are available within the manuscript and Supplementary files.

## Author contributions

G.W. and H.T. conceived the idea. H.T. and J.C. obtained funding for this research. L.L. and J.C. proposed and tested chicken breast model to study phototoxicity. G.W., J.E.S., and H.T. conducted related experiments. L.L., G.W., J.E.S., and H.T. performed data analysis and drafted the manuscript. H.T. and J.C. reviewed and edited the manuscript with inputs from all authors.

## Acknowledgments

The authors thank Stephen A. Boppart for sharing his laboratory, thank Rishyashring R. Iyer for his technical support, and thank Edita Aksamitiene, Eric J. Chaney, and Jennifer Mitchell for preparing biological samples. H.T. acknowledges the financial support from the National Institutes of Health, U.S. Department of Health and Human Services (R01 CA241618). J.C. acknowledges the financial support from the National Natural Science Foundation of China (82171991), the Special Funds of the Central Government Guiding Local Science and Technology Development (2020L3008), Fujian Major Scientific and Technological Special Project for “Social Development” (2020YZ016002) and the Natural Science Foundation of Fujian Province (2022J01216). J.E.S. was supported by NIBIB/NIH under Award Number T32EB019944.

---

**Supplementary Table 1.**
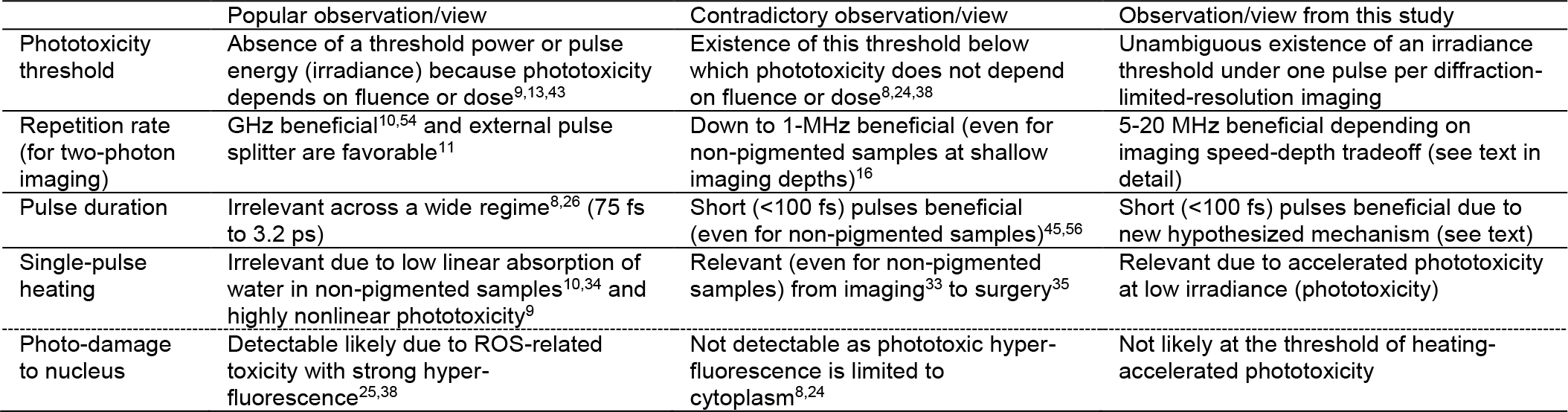
Apparent contradictions in gentle laser-scanning nonlinear optical imaging.

**Supplementary Table 2.**
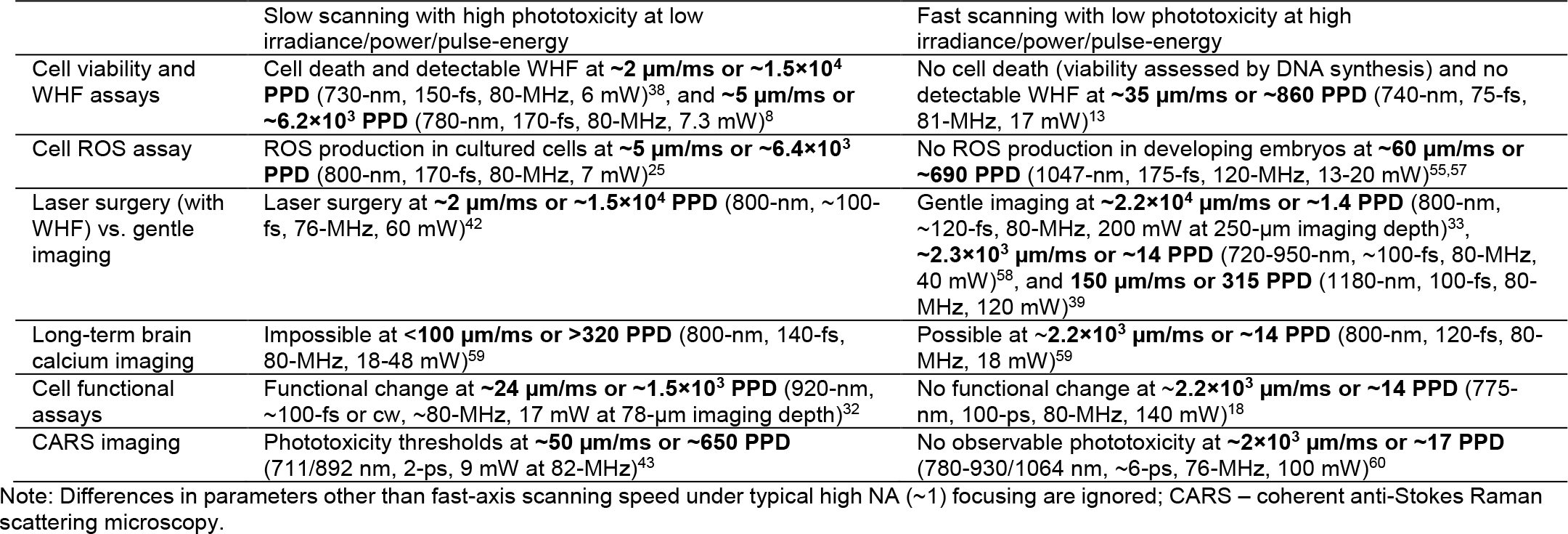
Dependence of NIR phototoxicity on fast-axis scanning speed or pulses per diffraction-limited-resolution (PPD) of ∼80-MHz pulses.

**Supplementary Table 3.** Noncomprehensive survey of label-free nonlinear optical imaging in biology and medicine.

| Modality/Application | Representative study or review |
| --- | --- |
| Second-harmonic generation microscopy | Campagnola, P.J. & Loew, L.M. Second-harmonic imaging microscopy for visualizing biomolecular arrays in cells, tissues and organisms. *Nat. Biotechnol.* **21**, 1356-1360 (2003). |
| Coherent Raman microscopy | Cheng, J.X. & Xie, X.S. Vibrational spectroscopic imaging of living systems: An emerging platform for biology and medicine. *Science* **350**, aaa8870 (2015). |
| Third-harmonic generation microscopy | Weigelin, B., Bakker, G.J. & Friedl, P. Third harmonic generation microscopy of cells and tissue organization. *J. Cell Sci.* **129**, 245-255 (2016). |
| Optical metabolic imaging | Kolenc, O.I. & Quinn, K.P. Evaluating cell metabolism through autofluorescence imaging of NAD(P)H and FAD. *Antioxid. Redox Signal.* **30**, 875-889 (2019). |
| Preclinical imaging | Dunn, K.W. et al. Functional studies of the kidney of living animals using multicolor two-photon microscopy. *Am. J. Physiol. Cell Physiol.* **283**, C905-16 (2002). |
| Developmental biology | Sun, C.K. et al. Higher harmonic generation microscopy for developmental biology. *J. Struct. Biol.* **147**, 19-30 (2004). |
| Dermatology | Chen, S.Y., Wu, H.Y. & Sun, C.K. In vivo harmonic generation biopsy of human skin. *J. Biomed. Opt.* **14**, 060505 (2009). |
| Drug discovery | Tipping, W.J., Lee, M., Serrels, A., Brunton, V.G. & Hulme, A.N. Stimulated Raman scattering microscopy: an emerging tool for drug discovery. *Chem. Soc. Rev.* **45**, 2075-2089 (2016). |
| Histology | Orringer, D.A. et al. Rapid intraoperative histology of unprocessed surgical specimens via fibre-laser-based stimulated Raman scattering microscopy. *Nat. Biomed. Eng.* **1**, 0027 (2017). |
| Cell therapy | Walsh, A.J. et al. Classification of T-cell activation via autofluorescence lifetime imaging. *Nat. Biomed. Eng.* **5**, 77-88 (2021). |
| Assisted reproductive technology | Shah, J.S., Venturas, M., Sanchez, T.H., Penzias, A.S., Needleman, D.J. & Sakkas, D. Fluorescence lifetime imaging microscopy (FLIM) detects differences in metabolic signatures between euploid and 20neuploidy human blastocysts. *Hum. Reprod.* **37**, 400-410 (2022). |
| Ophthalmology | Boguslawski, J. et al. In vivo imaging of the human eye using a 2-photon-excited fluorescence scanning laser ophthalmoscope. *J. Clin. Invest.* **132**, e154218 (2022). |

**Supplementary Table 4.** Reinterpretation of observed nonlinear NIR phototoxicity in unlabeled non-pigmented samples.

|  | Evidence supporting multiphoton-absorption-mediated phototoxicity | Reinterpretation to reconcile with linear-absorption-mediated nonlinear phototoxicity |
| --- | --- | --- |
| Power laws in cell assays (0.7-1.0 $\mu\text{m}$ excitation) | Power- and/or pulse-duration-dependent phototoxic assays attain an apparent multi-photon ($\geq 2$) order<sup>8,9,26</sup> | Acceleration by photoionization and heating of an otherwise linear phototoxicity attains apparently nonlinear phototoxicity |
| Pulsed versus cw phototoxicity | Absence of phototoxicity when pulsed excitation is switched to cw excitation (with larger powers)<sup>8,61,62</sup> | Unlike linear phototoxicity<sup>32,41</sup>, linear-absorption-mediated nonlinear phototoxicity accelerates with increasing power |
| Linked auto-fluorescence bleaching and lesion | Coincidence of photon order in bleaching and lesion generation ($\sim 4$) suggests NADH as a photosensitizer<sup>63</sup> | Acceleration by photoionization-heating and poor indication of phototoxicity by photo-bleaching result in this effect |
| Cellular ROS production and metabolic change (0.7-1.0 $\mu\text{m}$ excitation) | Similarity to UV-visible-induced ROS and metabolic change suggests multiphoton (rather than single-photon) excitation of intrinsic UV-visible sensitizers<sup>25,38</sup> | Single-photon excitation of NIR intrinsic photosensitizers with efficient intersystem crossing to triplet state induces this ROS production and metabolic change alternatively |
| Confined phototoxicity in one illumination plane (1.0-1.2 $\mu\text{m}$ excitation) | More phototoxicity from single-plane illumination over multiplane illumination of developing embryos suggests a multiphoton origin<sup>39</sup> | Linear-absorption-mediated nonlinear phototoxicity attain an apparent nonlinearity to confine phototoxicity in one illumination plane |
| Water absorption in a high NA microscope objective | Low and wavelength-dependent water absorption prohibits related heating as source for phototoxicity<sup>10,33</sup> | Exclusion of heating via water as the primary source for phototoxicity does not rule out linear phototoxicity from a non-water NIR photosensitizer in the hypothesized mechanism |
| Nonlinear fluorescence photo-bleaching | Fluorescence bleaching attain a $>2$ photon order (higher than that of "photo-enhancement")<sup>64,65</sup> | Photo-bleaching of intrinsic and extrinsic fluorophores is a poor indicator of phototoxicity<sup>2,3</sup> |

**Supplementary Table 5.**
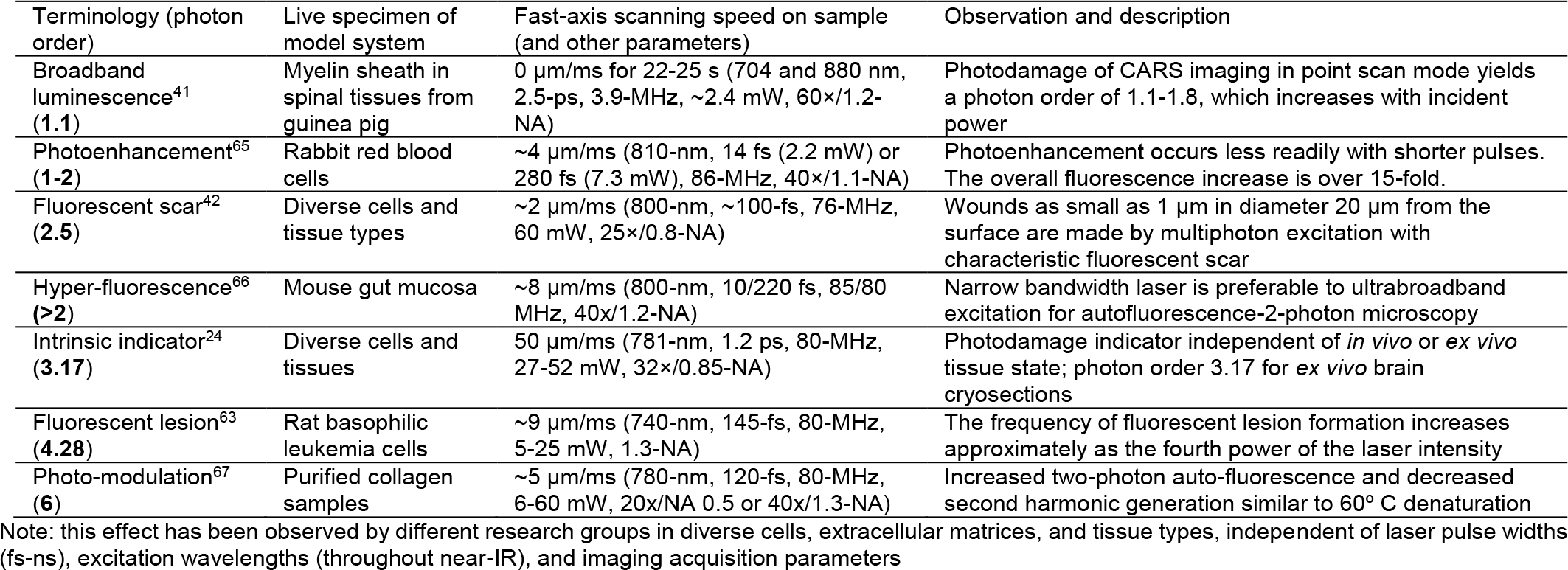
Hyper-fluorescence-like effects observed with diverse multiphoton illuminations in unlabeled non-pigmented live specimens.

**Supplementary Table 6.** Quantitative phototoxicity bioassays with multiphoton excitation.

| Nature of bioassay (photon order) | Live specimen of model system | Fast axis scanning speed on sample (and typical other parameters) | Details of bioassay |
| --- | --- | --- | --- |
| Inline labeled<sup>32</sup> (1) | Cortical astrocytes in mouse brain slices | $\sim 24$ $\mu\text{m}/\text{ms}$ (920-nm, $\sim 100$-fs or cw, $\sim 80$-MHz, 17 mW, 20x/0.95-NA) | Calcium microdomain hyperactivity |
| Offline labeled<sup>68</sup> (1.19-1.28) | Drosophila melanogaster | 0 $\mu\text{m}/\text{ms}$ (800-nm, 37/100-fs, 1-kHz, irradiance 0.1 TW/cm<sup>2</sup>, no focusing) | TUNEL cell assay in salivary glands superficially located in the larva's body |
| Inline labeled<sup>26</sup> (2) | Neocortical neurons in rat brain slices | $\sim 2$ $\mu\text{m}/\text{ms}$ (870-nm, 75-fs with 3-7 mW or 3.2-ps with 8-24 mW, $\sim 80$-MHz, 60x/0.91-NA) | Basal fluorescence of various Ca<sup>2+</sup>-indicators |
| Offline label-free<sup>8</sup> (2) | Chinese hamster ovary cells | $\sim 5$ $\mu\text{m}/\text{ms}$ (780-nm, 170-fs, 80-MHz, 7.3 mW, 40x/1.3-NA) | Reduced cloning efficiency (clone consists of $<8$ cells) |
| Inline labeled and offline label-free<sup>9</sup> (2.5) | Bovine adrenal chromaffin cells | 25 $\mu\text{m}/\text{ms}$ (840-nm, 190-fs, 82-MHz, 7-20 mW, 63x/0.9-NA) | Changes in resting [Ca<sup>2+</sup>] level via FURA-2 and degranulation reaction |
| Offline label-free<sup>39</sup> (2-3) | Drosophila embryos | 150 $\mu\text{m}/\text{ms}$ (1180-nm, 100-fs, 80-MHz, 120 mW, 20x/0.95-NA) | Embryo survival rate and cellularization speed |
| Offline labeled<sup>69</sup> (2-3) | Chinese hamster ovary cells | 8-20 $\mu\text{m}/\text{ms}$ (695-810 nm, 130-fs, 80-MHz, 14 mW typically, 40x/0.8-NA) | Damage to nucleus like that from exposure to solar ultraviolet light |

**Supplementary Video 1**. Time-lapse eSLAM imaging of hamster kidney cells showing heterogeneous WHF (arrowhead in Extended Data Fig. 6b).

**Supplementary Video 2**. Time-lapse eSLAM imaging of *ex vivo* mouse kidney tissue showing heterogeneous WHF (arrowhead in Extended Data Fig. 6c).

**Supplementary Video 3**. Time-lapse pSLAM imaging of chicken breast showing early homogeneous WHF versus late heterogeneous WHF and bubble formation (SHG - green; 3PAF - cyan; 2PAF - yellow).

**Supplementary Video 4**. Time-lapse eSLAM imaging of *ex vivo* mouse brain slice showing homogeneous WHF via 3PAF (cyan) but not 2PAF (yellow) or THG (magenta).

**Supplementary Video 5**. Time-lapse eSLAM imaging of moving *C. elegans* worms.

